# Multiscale analysis of myelin alterations in skin biopsies from synucleinopathies

**DOI:** 10.1101/2025.05.15.654285

**Authors:** Marta Di Fabrizio, Bram L. van der Gaag, Mara Terzi, Eve Aaron, Noa B. C. Steerenberg, John G. J. M. Bol, Wilma D. J. Van de Berg, Henning Stahlberg, Amanda J. Lewis

## Abstract

Loss of myelin and demyelination play a role in the pathophysiology of Parkinson’s disease (PD) and related neurodegenerative diseases, but little is known about the ultrastructure of the myelin-axon unit in the peripheral nervous system of subjects diseased with synucleinopathies. We here present an analysis of the myelin ultrastructure and the myelin protein abundance that characterize myelinated axons of nerve fiber bundles in cervical skin biopsies of 45 pathologically confirmed PD, DLB, MSA and non-neurological control donors. We calculated a myelin damage score and classified over 1100 myelin sheaths by looking at myelin fragmentation and swellings with correlative light and electron microscopy. We found a higher load of myelin damage in the PD compared to MSA and control groups. Quantification with ELISA did not reveal any differences in myelin protein zero (MPZ) concentrations in skin tissue homogenates between synucleinopathies. The observed structural abnormalities in the myelin sheaths may help to discriminate among subjects with synucleinopathies and control subjects and to understand the involvement of the peripheral innervation in the diseases. Our multiscale analysis of peripheral nerves highlights their potential as future biomarkers for the detection and differentiation of synuclein diseases.

## Introduction

Synucleinopathies (Parkinson’s disease – PD, Dementia with Lewy bodies – DLB and multiple system atrophy – MSA) are neurodegenerative diseases characterized by the presence of misfolded alpha-synuclein (aSyn) aggregates in the brain^1–4^. Nevertheless, it is still a matter of debate whether these aggregated aSyn species are the cause or the consequence of synucleinopathies, as other factors contribute to pathogenesis such as oxidative stress, proteasomal and mitochondrial dysfunction^5–8^, neuroinflammation and impaired cellular trafficking pathways^9,10^ and myelin alterations^11–16^. Macroscopic-scale imaging and omics studies on myelin alterations in the central nervous system have shown axonal and myelin abnormalities to play a crucial role in the development of synucleinopathies^11–17^ and other neurodegenerative diseases^18–21^.

A complex interplay between multiple factors seems to lead to the development of synucleinopathies^22,23^. In recent years, great efforts were made in characterizing the presence of aSyn deposits in the nerve fibers in peripheral tissues to unravel their role in the etiology and progression of these diseases. aSyn pathology and seeding propensity measured in skin biopsies show strong potential as a diagnostic tool for synucleinopathies^24–32^ since the tissue is easily accessible and collection procedures are well-tolerated. Both, PD and DLB subjects show signs of peripheral neuropathy in up to 55% of the cases^33,34^. In these studies, neuropathy was characterized as a reduction of nerve fiber density and/or conduction speed of action potentials along the axons. Thus, myelination, which forms the lipid membrane around neuronal axons allowing rapid saltatory conduction^20,35^, might also be affected in the peripheral tissues of these diseased subjects.

These findings suggest axonal damage and demyelination in the central and peripheral nervous systems in synucleinopathies, but little is known about if and how myelin sheaths in the periphery are impaired. Changes in the structure of myelin sheaths might provide insights in the pathophysiology underlying autonomic and motor symptoms in these disorders. Novel approaches and strategies are needed to identify peripheral markers for synucleinopathies.

In this study, we provide detailed insight into the myelin ultrastructure and myelin protein abundance that characterizes myelinated axons of nerve fiber bundles in cervical skin biopsies from 45 pathologically confirmed PD, DLB, MSA and non-neurological control donors, using correlative light and electron microscopy (CLEM). We combined this technique with an in-house developed enzyme linked immunosorbent assay (ELISA) to quantify levels of myelin protein zero (MPZ), responsible for myelin sheath compaction in the peripheral nervous system^20,36^. Finally, we showed that myelin damage correlates with the presence of hallucinations and advanced aSyn pathological stages. Our results provide strong evidence for a differential mechanism in myelin preservation in synucleinopathies that shows potential as a biomarker for disease differentiation.

## Results

### The ultrastructure of myelin sheaths in skin biopsies from aged individuals

We performed an ultrastructural assessment to study the integrity of the myelin sheaths in the human dermis, collected at the cervical region C6-7, from 36 deceased subjects (16 PD, 4 DLB, 5 MSA and 11 non-neurological controls). The demographics, clinical data and neuropathological staging are shown in Supplementary Tables 1 and 3.

Using CLEM, we identified by light microscopy myelinated nerve fiber bundles in the human dermis on 200 nm consecutive sections with antibody labelling against MPZ and toluidine blue staining (Figure 1a and b). With transmission electron microscopy (TEM), we imaged the TEM grids at the correlated locations on sections adjacent to those on the glass slides (Figure 1c and d), analyzing 1123 myelin sheaths in total (384, 465, 104 and 170 from controls, PD, DLB and MSA, respectively).

**Figure 1:**
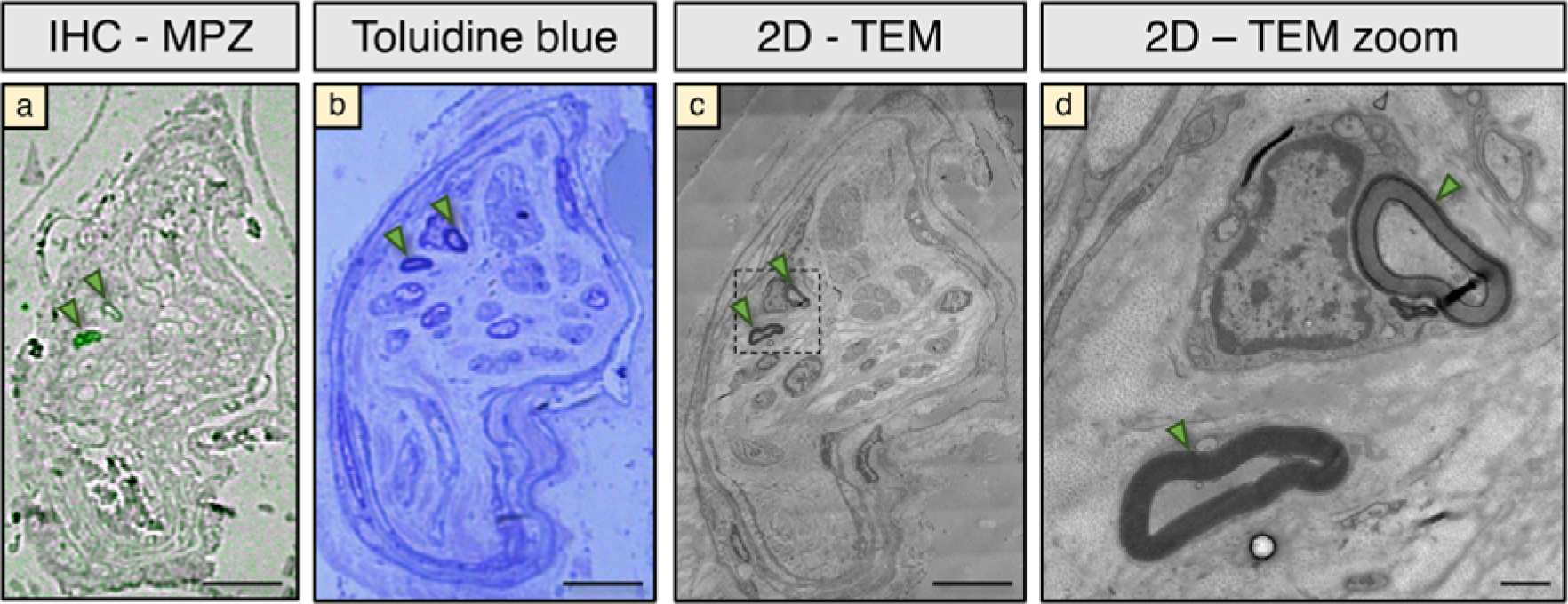
Localization of nerve fiber bundles in the human dermis using CLEM. (a) and (b) consecutive 200 nm sections stained with immunohistochemistry (IHC) against MPZ and toluidine blue, correlated with (c) adjacent 80 nm section containing the same nerve fiber bundle imaged with TEM and (d) higher magnification. Arrowheads indicate myelin sheaths. Scale bars a, b, c: 10 µm, d: 1 µm.

To assess the myelin sheath integrity, we defined 4 categories of myelin damage (absent N-, mild N+, moderate N++ and high damage N+++) based on the size and density of myelin sheath swellings visible with TEM (Figure 2a-h, Supplementary Figures 1-2). The criteria used to classify the myelinated fibers are discussed in the Methods. In total, we found N-=334, N+=555, N++=197 and N+++=37 myelinated fibers, from at least 18 myelinated axons per donor. Electron tomography was performed on selected myelin sheaths to capture the 3D arrangement of compact and swollen myelin (Supplementary Figure 3).

**Figure 2:**
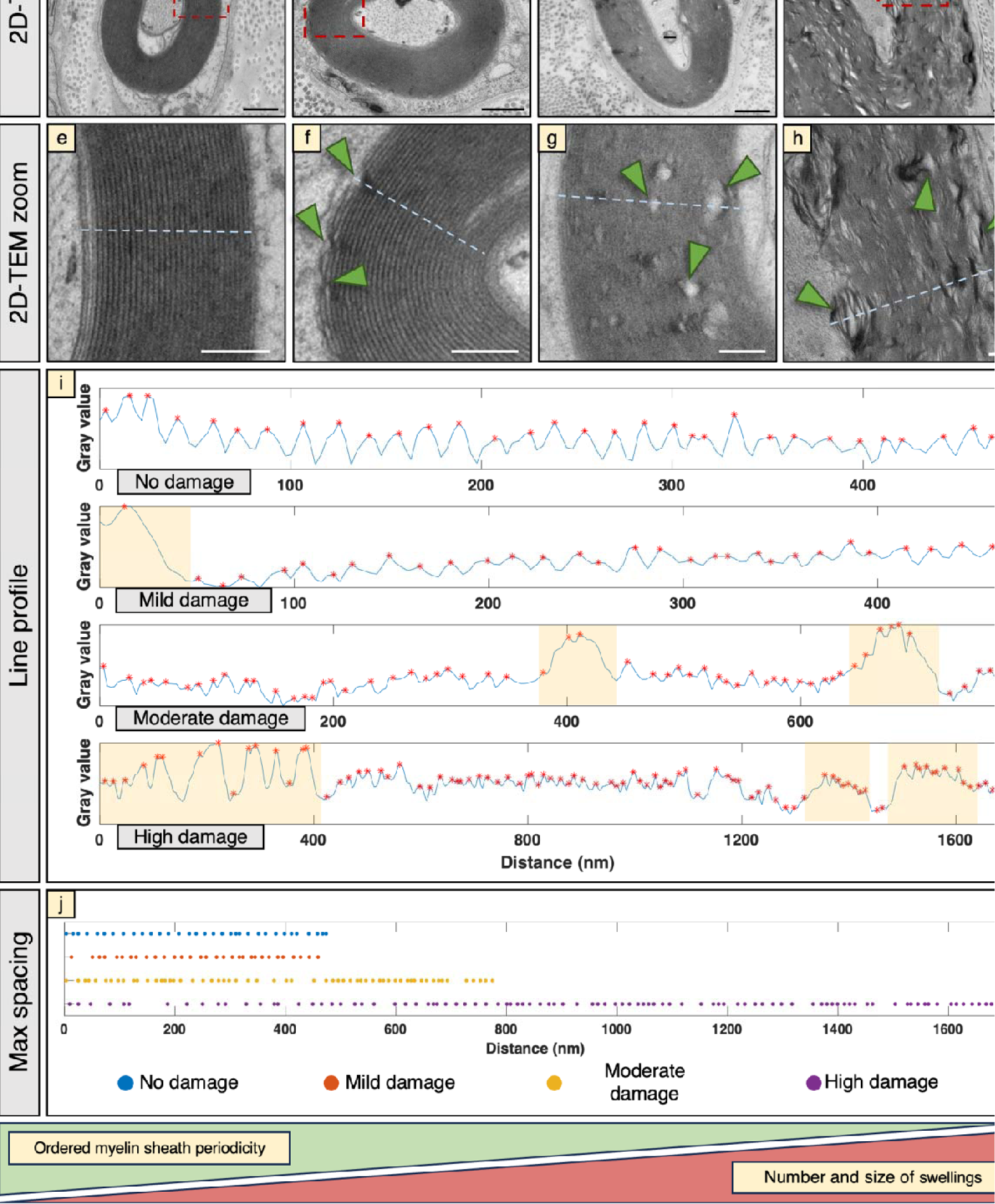
Classification of myelin damage. (a) - (d) Four categories of myelin damage was identified based on 2D – TEM ranging from no (N-), mild (N+), moderate (N++) and high (N+++) damage. (e) - (h) Zoomed red dashed squares from (a) – (d) showing myelin sheath ultrastructure in close detail. Examples of myelin sheath swellings are indicated with arrowheads. Ranging from no to high damage, the myelin sheath arrangement becomes less ordered as the number and size of swelling increase. (i) Examples of line profiles (light blue) for the four damage categories, the red asterisks indicate the local maxima in the line profiles. Light blue dashed lines in (e) – (h) indicate the location in the TEM image the line profile was taken from. Light yellow bands indicate the widest peaks in the line profiles, corresponding to swellings in the myelin sheaths. (j) Local maxima spacing in the four categories. Line profiles from no and mild damaged images show high periodicity with local maxima almost equally spaced along the distance. Due to the increasing number of swellings, the periodicity in moderately and highly damaged myelinated fibers is reduced. Moreover, the width of the peaks in the line profile is constant in the case of no and mild damage, while is quite irregular for moderate and high damage. Scale bars (a) – (d): 500 nm, (e) – (h): 200 nm

### Disease groups show differences in myelin preservation

Based on the classification of myelin damage shown in Figure 2, we calculated the proportion of damaged myelin sheaths in the four groups (controls, PD, DLB and MSA). Controls and MSA showed a higher proportion of preserved (undamaged) myelin sheaths, approximately 40%, compared to about 20% in PD and DLB (Figure 3a). The proportion of mildly damaged myelin was relatively constant across the four groups (about 50%), while PD and DLB showed a higher fraction of moderately and highly damaged myelin sheaths compared to controls and MSA. A chi-squared test confirmed significant differences in the distribution of damage categories between groups (Figure 3b). Standardized residuals (R) indicated an increased number of preserved sheaths in controls and MSA, and a corresponding decreased number in PD and DLB. Conversely PD and DLB showed increased moderate and severe damage relative to controls and MSA, highlighting a disease-specific pattern of myelin pathology.

**Figure 3:**
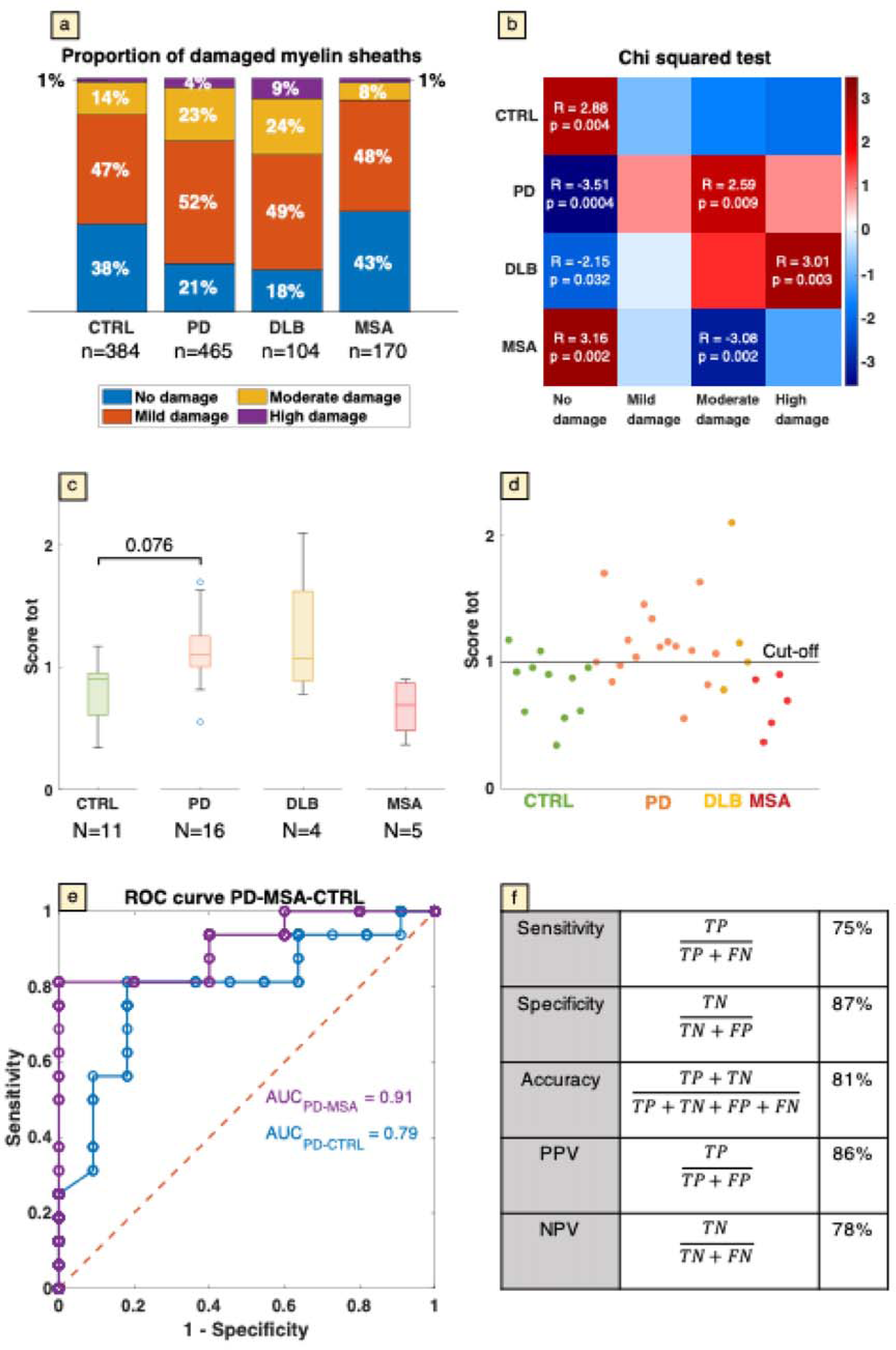
Disease groups show differences in myelin preservation. (a) Proportion of the four myelin damage classes in each group. n indicates the number of myelin sheaths analyzed per category. (b) Chi-squared test showing standardized residuals (R) and p-values corrected for multiple comparisons according to the Bonferroni method. Only significant p-values are shown. Both in (a) and (b) CTRL and MSA show more preserved and less moderately and highly damaged myelin sheaths, while PD and DLB show the opposite. (c) Box and whisker plot showing PD group with significantly higher levels of myelin damage compared to non-neurological control groups (p<0.05). Empty circles are outliers calculated with the standard interquartile range method. N indicates the number of donors in each category. The horizontal line in the box plots represents the median. Significant p-values are shown. The p-value between control and PD was calculated with a two-sample t-test. DLB and MSA were not considered for the p-value calculation due to the small sample size N. (d) Total score plot showing the individual donor differences across the disease groups. PD group shows higher total score compared to non-neurological control and MSA groups. The cut-off line at score tot = 1 is used to discriminate among high (>1) and low (<1) myelin damage classes. (e) ROC curve showing an area under the curve (AUC) of 0.91 and 0.79 for PD-MSA and PD-CTRL respectively. The red dashed line represents the random guess. (f) Summary table with calculated values for sensitivity, specificity, accuracy, positive predictive value (PPV) and negative predictive value (NPV). TP = true positive, TN = true negative, FP = false positive, FN = false negative.

We next defined a total score for each donor as a weighted mean of the four categories of myelin damage, by assigning score values of 0, 1, 2, or 4, to completely preserved, mildly, moderately and highly damaged myelin sheaths, respectively, using the *total score* formula in the Methods. We investigated if the total score of myelin damage was an effective metric to discriminate among disease groups. We found differences in the load of myelin sheath preservation and myelin wrapping compaction in the nerve fiber bundles from the skin of PD compared to MSA and controls (Figure 3c). More specifically, the PD group showed a significantly higher total score for myelin damage compared to control (p-value: 0.0076). A similar trend was observed between PD and MSA groups. However, as the small sample size of the DLB and MSA groups (N=4 and N=5, respectively) limited statistical inference, we excluded them from the statistical analysis. We therefore report descriptive statistics for the myelin damage as mean ± SEM for each group: CTRL 0.82 ± 0.08, PD 1.13 ± 0.07, DLB 1.26 ± 0.29, MSA 0.67 ± 0.10. This suggests a higher myelin damage load in PD and DLB compared to controls and MSA. No significant difference was found between control and MSA groups. To assess group differences while controlling for age at death, we applied an analysis of covariance (ANCOVA) and found that age had no significant influence on the total score (p=0.46; data not shown).

Inspecting the total scores within each disease group, we found that 75% of PD donors (12/16), and 75% (3/4) of the DLB groups showed a total score equal or greater than one, indicating that they exhibited extensive myelin damage, with the greatest contribution coming from the moderate and high myelin damage categories (Figure 3d). Conversely, 82% (9/11) of control donors and 100% (5/5) of MSA donors showed a myelin damage total score smaller than one, indicating that they exhibited little myelin sheath damage, where no damage and mild damage categories had the greatest contribution. We therefore suggest that the level of myelin damage could serve as classifier to differentiate between synuclein diseases, using a cut-off for the total score of 1.

In addition, we calculated the receiver operating characteristic curve (ROC) and the area under the curve (AUC) to evaluate the performance of the proposed classifier in discriminating between PD/controls and PD/MSA. The AUC was 0.91 in the case of PD-MSA and 0.79 for PD-CTRL, suggesting that there was a 91% and 79% chance of distinguishing PD from MSA and controls, respectively (Figure 3e). Given the small number of DLB donors and variability in their scores - 50% (2/4) had a score > 1, 25% (1/4) had a score < 1 and 25 % had a score = 1-no conclusions could be drawn for disease group. Therefore, DLB cases were excluded from the sensitivity, accuracy, and predictive value calculations. For PD, CNTRL and MSA, we report a sensitivity of 75% (12/16), a specificity of 87% (14/16), an accuracy of 81% (26/32), a positive predictive value (PPV) of 86% (12/14) and a negative predictive value (NPV) of 78% (14/18), suggesting an overall good performance of this classifier (Figure 3f).

### Myelin thickness and g-ratio remain constant across disease groups, but show differences according to the level of damage

TEM studies of myelin routinely assess thickness and g-ratio of myelinated axons as they are important parameters for understanding the efficiency and function of myelinated nerve fibers in the nervous system^37^. Thicker myelin generally corresponds to faster nerve signal transmission. The g-ratio is a measure of how thick the myelin sheath is with respect to the axon size. Typical g-ratios in the PNS vary between 0.6 and 0.7^37^. Higher g-ratios indicate thinner myelin, often seen in demyelinating diseases, while lower g-ratios indicate excess myelin. Diseases that alter myelin affect both parameters, impacting nervous system function.

We therefore quantified myelin thickness and g-ratio across disease groups and according to the level of damage (absent, mild, moderate and high). No significant differences were found in myelin thickness and g-ratio between the disease groups, except for a slight decrease in the myelin g-ratio for DLB donors compared to controls (Figure 4a, c). However, when looking at the different categories of myelin damage, the non- and mildly damaged myelin sheaths were significantly thinner compared to the moderately and highly damaged sheaths (Figure 4b) as a result of absence or low amount of swellings in these categories. Similarly, the g-ratio was significantly lower for highly damaged myelin sheaths (Figure 4d), in accordance with increased myelin thickness. Analysis with ANCOVA showed no influence of age in any of the panels (a: p=0.12, b: p=0.055, c: p=0.42, d: p=0.076).

**Figure 4.**
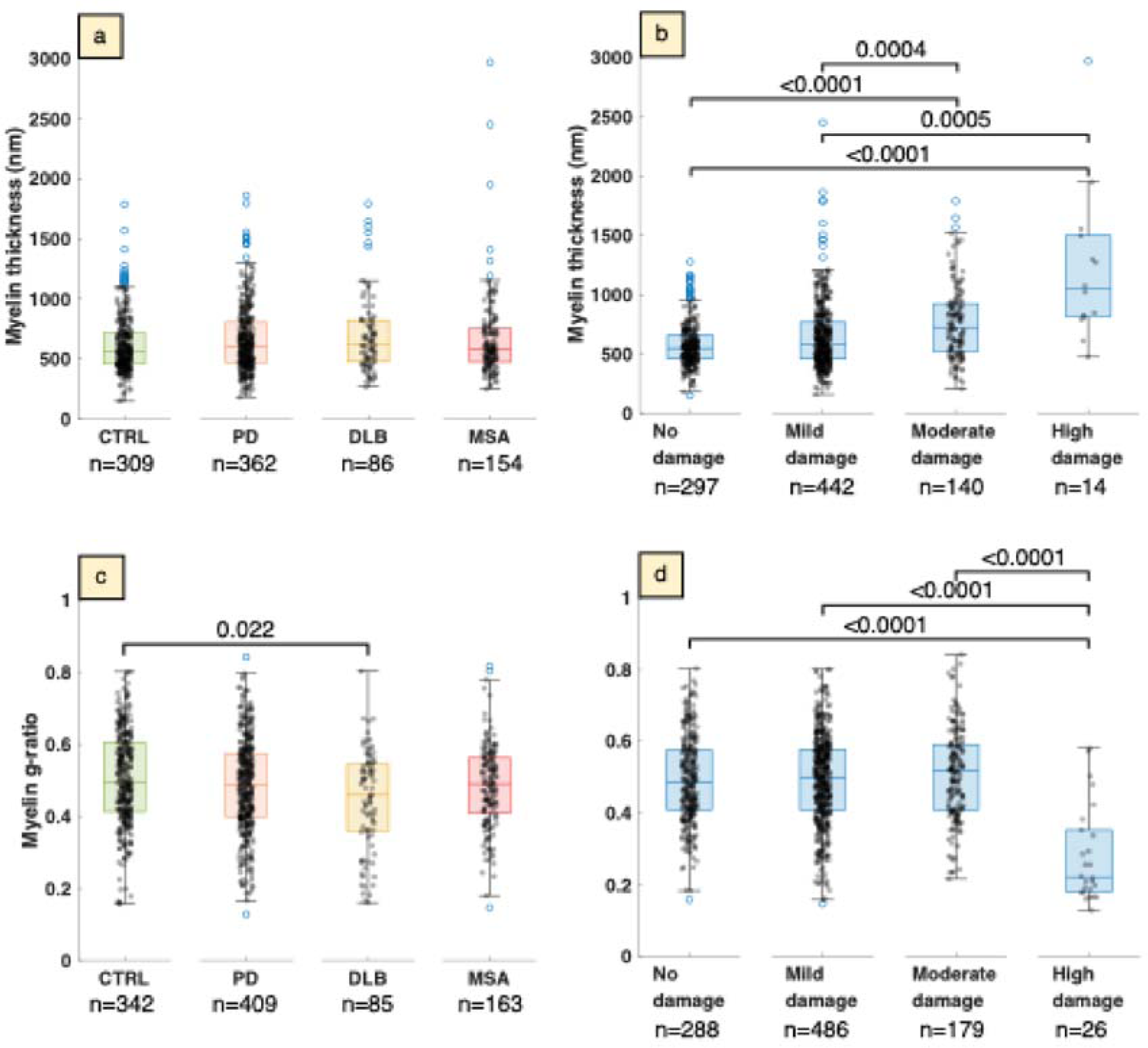
: Myelin thickness and g-ratio across disease groups and damage categories. Box and whisker plot showing myelin thickness and g-ratio. Thickness (a) and g-ratio (c) did not show any significant difference in the cohort except for a slight decrease in g-ratio when comparing DLB and control groups. (b) However, myelin sheaths were significantly thicker in moderate and highly damaged categories. (d) g-ratios for no, mild and moderate damage categories were significantly increased compared to the highly damaged group. P-values were calculated with Kruskal-Wallis test followed by Dunn’s test and corrected for multiple comparisons according to the Bonferroni method. Only significant differences (p<0.05) are indicated (brackets). The horizontal line in the box plots represents the median. Empty circles are outliers calculated with the standard interquartile range method. n is the number of myelin sheaths analyzed per group.

### Myelin damage is not affected by differences in post-mortem delay, fixation time or age at death

In our cohort, all skin biopsies were collected at autopsy with the same standardized protocol and with a post-mortem delay (PMD) below 10 hours. Since aging, PMD and fixation are common factors known to affect myelin^38,39^ and tissue^40^ integrity, we assessed if any of these factors correlated with the level of myelin damage observed in our samples. First, we assessed the overall relationship between these factors and myelin damage across the entire cohort, regardless of the disease group. To do this, the total myelin damage score for each individual donor was plotted as a function of either PMD, fixation time or age at death. No significant correlation was observed (correlation coefficients c = -0.07, -0.23 and -0.04, respectively, and p-values > 0.05; Figure 5a, b and Supplementary Figure 5a).

**Figure 5:**
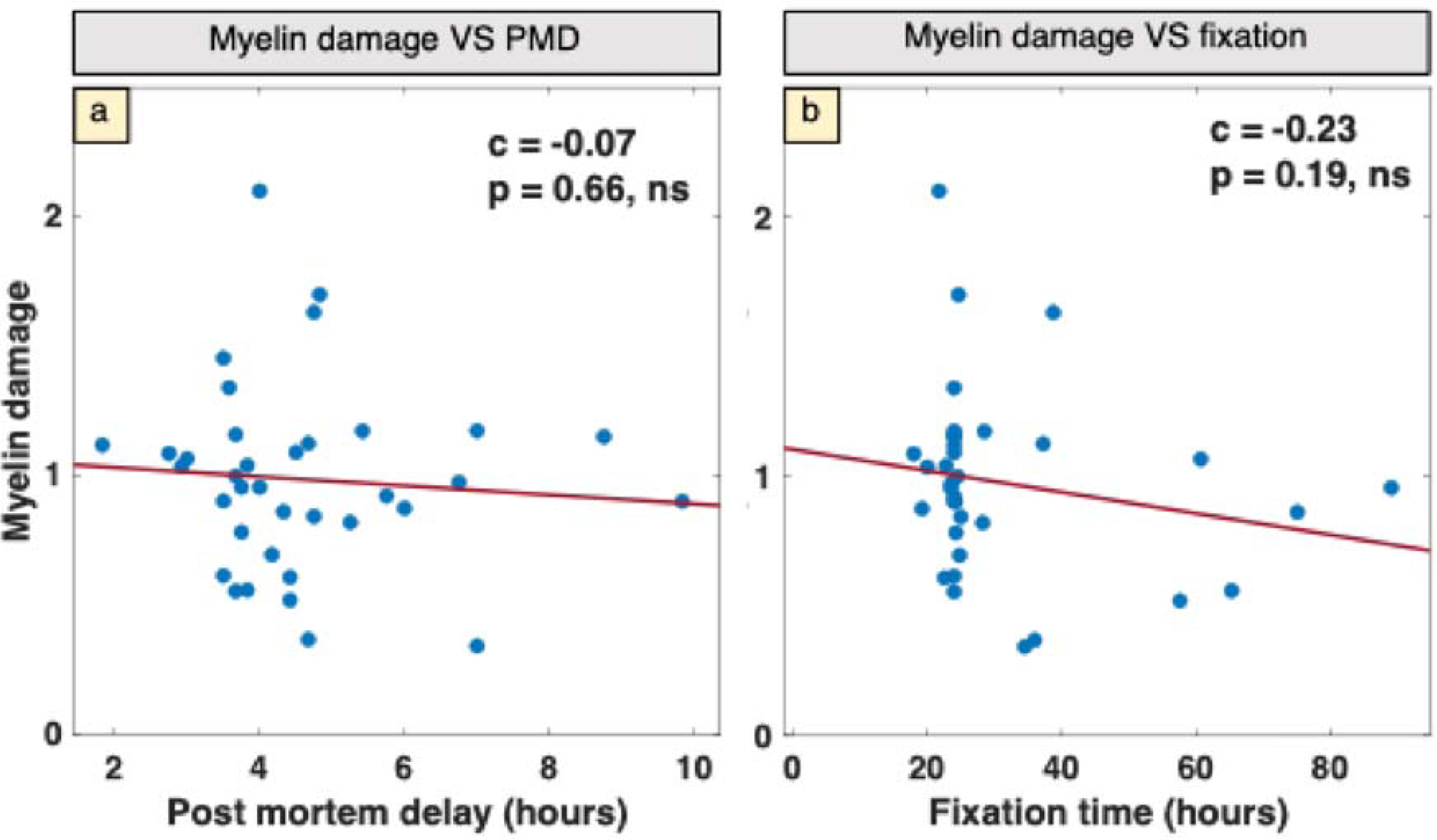
The effect of post-mortem delay (PMD) and fixation time on the differences in myelin damage. Correlation plots of total myelin damage score as a function of (a) PMD and (b) biopsy fixation time at autopsy. No significant correlation (p>0.05) was detected between PMD/fixation time and myelin damage. c = Spearman correlation coefficient, ns = non-significant.

To determine whether the absence of correlation held true within individual disease categories, we next correlated the same relationships within each diagnostic group, specifically, controls and PD. Again, no significant correlations were found between PMD, fixation time, age and myelin damage in either group (Supplementary Figures 4, 5).

The DLB and MSA groups were excluded from this sub-group analysis due to the small sample size (N=4 and N=5, respectively), which limit statistical power. However, a strong positive correlation between myelin damage and age at death was observed in the MSA group (c=0.98, p=0.004). Caution is warranted in interpreting this result, as small sample sizes can inflate correlation coefficients and yield misleading statistical significance. Notably, the MSA group had a significantly younger average age at death than for all other groups (Supplementary Table 1) likely due to an earlier age of onset and shorter disease duration. While 3 out of 5 MSA cases in our cohort died by euthanasia, this was also the case for several PD patients (Supplementary Table 3). Importantly, even when accounting for euthanasia, MSA is widely recognized as a more aggressive disorder with a naturally shorter disease course compared to PD^41,42^. Despite the potential influence of age within the MSA group, their myelin damage levels remained comparable to the older control group, and remained well below those observed in the PD group, which had a similar age distribution to the control group (Figure 3c,d).

In summary, our findings indicate that differences in myelin damage among the disease groups in our cohort were not a consequence of PMD, fixation time, or age at death.

### Dermal myelin damage correlates with hallucinations and aSyn brain pathology

We next investigated how clinico-pathological data such as age at disease onset, disease duration, brain pathology, the presence of hallucinations, cognitive impairment (calculated as clinical dementia rating - CDR^43^) correlated with the extent of myelin damage observed by electron microscopy. These relationships were analyzed across the entire cohort, regardless of disease group, and are summarized as a correlation matrix (Figure 6).

**Figure 6:**
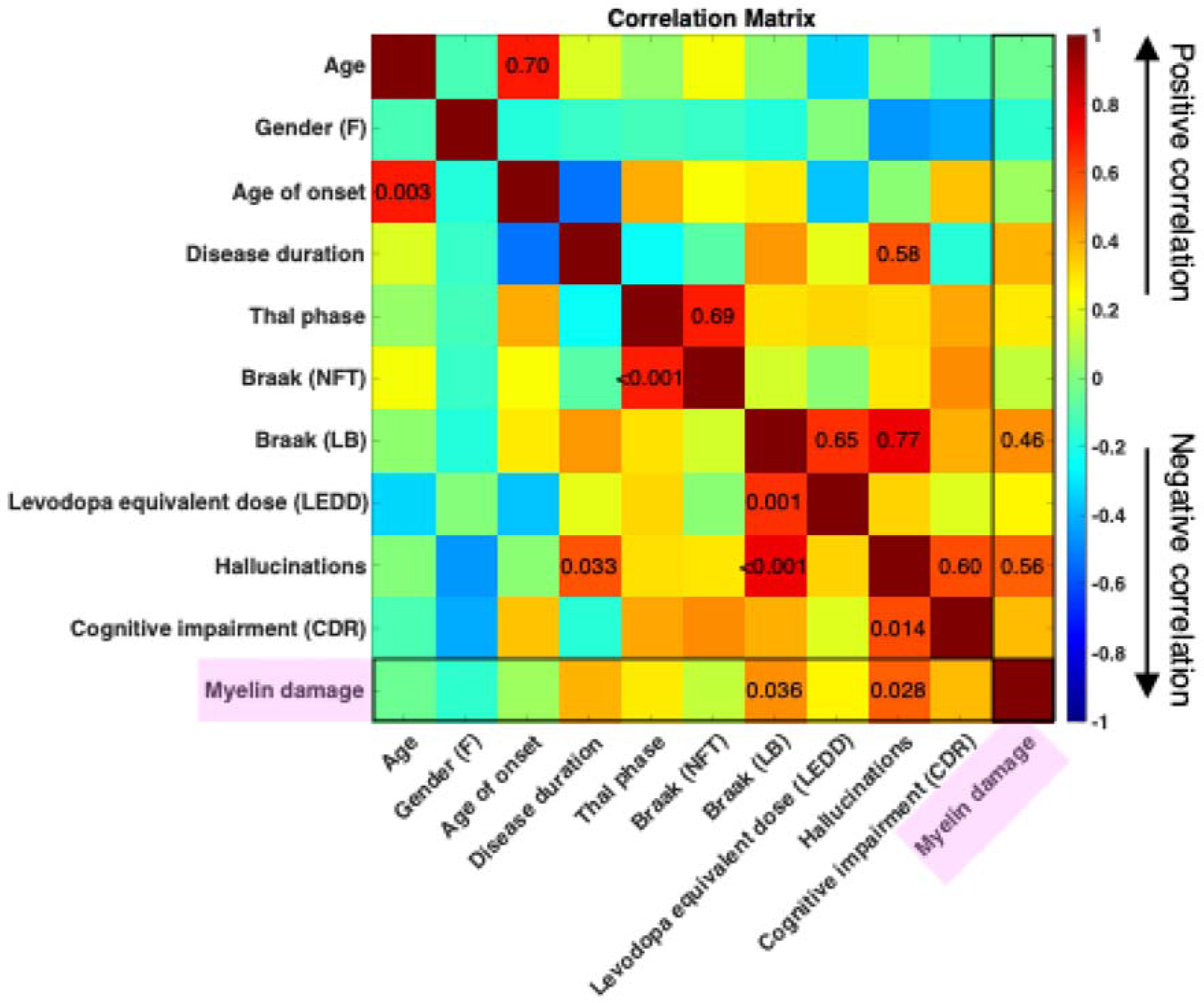
Correlation matrix of clinical symptoms and brain pathology with each other and with myelin damage. The values in the fields in the upper triangle are correlation coefficients, the values in the lower triangle are the p-values. Numbers are only shown for fields with significant correlation values (p<0.05). Warm and cold colors represent positive and negative correlations. Myelin damage (black boxes) shows correlation with the presence of hallucinations and high values of LB Braak staging. Binary variables: gender and hallucinations; ordinal variables: Thal, Braak (NFT), Braak (LB) and cognitive impairment; continuous variables: age, age of onset, disease duration, levodopa equivalent dose, myelin damage. The following correlation coefficients were calculated according to variable type and distribution: binary vs binary – Phi coefficient (equivalent to Pearson); binary vs continuous – point biserial coefficient (equivalent to Pearson); binary vs ordinal, ordinal vs ordinal, ordinal vs continuous, continuous vs continuous – Spearman coefficient. P-values were adjusted for multiple comparisons with the Benjamini-Hochberg procedure. F: female, NFT: Neurofibrillary tangle, LB: Lewy body.

As expected, the correlation matrix captured known relationships that serve as internal validation of the analysis. For example, there is a positive correlation between Thal phase vs Braak NFT stage and the presence of hallucinations vs cognitive impairment.

Focusing on myelin damage, the strongest correlation was shown to be with the presence of hallucinations (c = 0.56, p-value = 0.028). This likely reflects a group-level association, as almost all PD and DLB donors noted hallucinations, and exhibited higher levels of myelin damage, while hallucinations were absent in MSA and control donors, which also showed lower myelin damage.

Additionally, a moderate positive correlation was observed between myelin damage and Braak LB stage (c = 0.46, p-value = 0.036). However, this too likely reflects group differences, as all PD donors had advanced aSyn pathology with Braak LB stages 5 or 6.

Moreover, it is important to note that myelin damage did not show any significant correlation with the administered dose of levodopa, calculated as levodopa equivalent daily dose (LEDD^44^), in the three months before death. Therefore, the absence of an association between levodopa dose and myelin damage in our cohort rules out the possibility that myelin damage is a direct effect of levodopa-induced peripheral neuropathy.

### Skin biopsy myelin protein zero levels do not differ across synucleinopathies and controls

MPZ is considered to be the most abundant myelin adhesion protein in the peripheral nervous system and it is believed to be responsible for myelin compaction^36^. To assess the potential of an involvement of MPZ in the myelin alterations observed in our cohort, we measured total MPZ levels in the skin samples across all groups compared to controls using an ELISA approach (34 donors in total). No differences could be observed between PD (mean MPZ = 1.389 ng/mL), DLB (mean MPZ = 1.366 ng/mL), MSA (mean MPZ = 1.532 ng/mL), and control cases (mean MPZ = 1.267 ng/mL) (Figure 7A). When grouping the synucleinopathies (mean MPZ = 1.406 ng/mL) and comparing these with the controls, we could also not detect any significant difference (Figure 7B). These data suggest that the observed myelin damages are not related to altered levels of MPZ.

**Figure 7.**
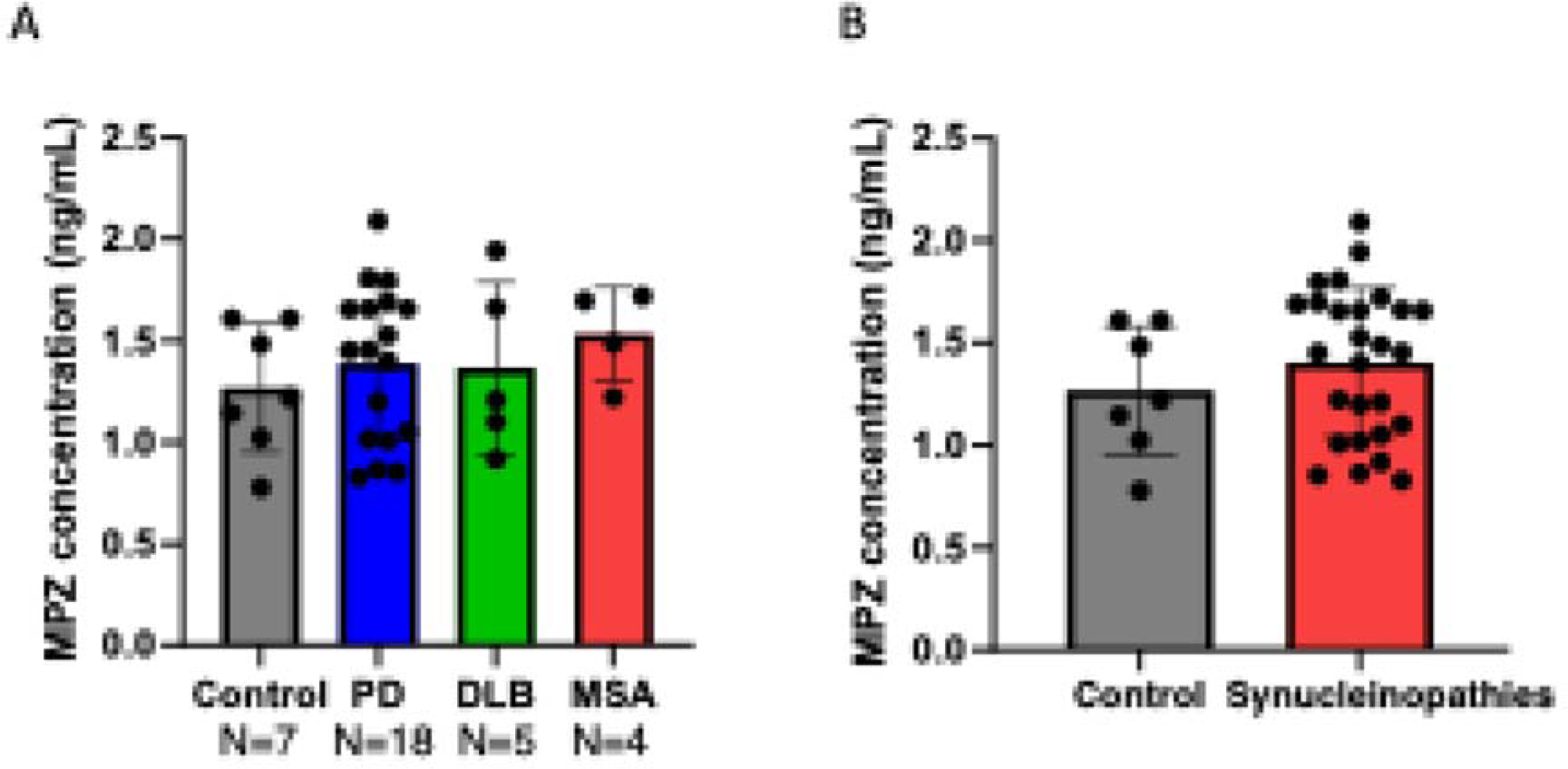
MPZ ELISA did not show any difference in MPZ concentration between synucleinopathies and controls. **(A)** MPZ values were corrected for total protein loading and did not show any differences between controls, PD, DLB and MSA cases (p>0.05). The p-values were calculated using a one-way ANOVA and corrected with Tukey’s test for multiple comparisons. (B) No differences were observed in corrected MPZ values between controls and synucleinopathies (p>0.05). The p-value was calculated using an unpaired t-test. Graphs display mean ± SD.

### The examined nerve fiber bundles are locally negative for alpha-synuclein aggregates

Given the association of myelin damage with LB pathology in the brain (Figure 6), we investigated if a similar association with myelin damage and peripheral aSyn pathology could also be found. Several studies show phosphorylated aSyn (pS129) accumulation in the dermal nerve fibers of patients and donors affected by synucleinopathies^24–31^. We therefore performed immunohistochemistry on paraffin sections from one PD case to assess the co-localization of MPZ with aSyn and PGP9.5 (pan-neuronal marker in the PNS) using confocal microscopy. pS129 aSyn deposits were detected in both non-myelinated (PGP9.5+/MPZ-) as well as in myelinated nerve fiber bundles (PGP9.5+/MPZ+) (Supplementary Figure 7).

We therefore assessed if the myelin damage observed with EM could be attributed to the local presence of pS129 aSyn in the same portions of nerve fiber bundles. We performed immunohistochemistry on selected 200 nm skin resin sections using a set of eight aSyn antibodies, with MSA brain tissue (*substantia nigra*) serving as a positive control for the primary antibodies (Supplementary Table 2). Toluidine blue staining of adjacent sections was used to confirm the tissue morphology and identify myelinated nerve fiber bundles in the skin. While all antibodies robustly labelled glial cytoplasmic inclusions in the MSA brain (example shown for clone EP1536Y in Figure 8a), none showed positive immunostaining in the skin nerve fiber bundles (orange dashed line in Figure 8b, c). These results suggest that the observed myelin damage in nerve fiber bundles was not associated with a local accumulation of aSyn.

**Figure 8.**
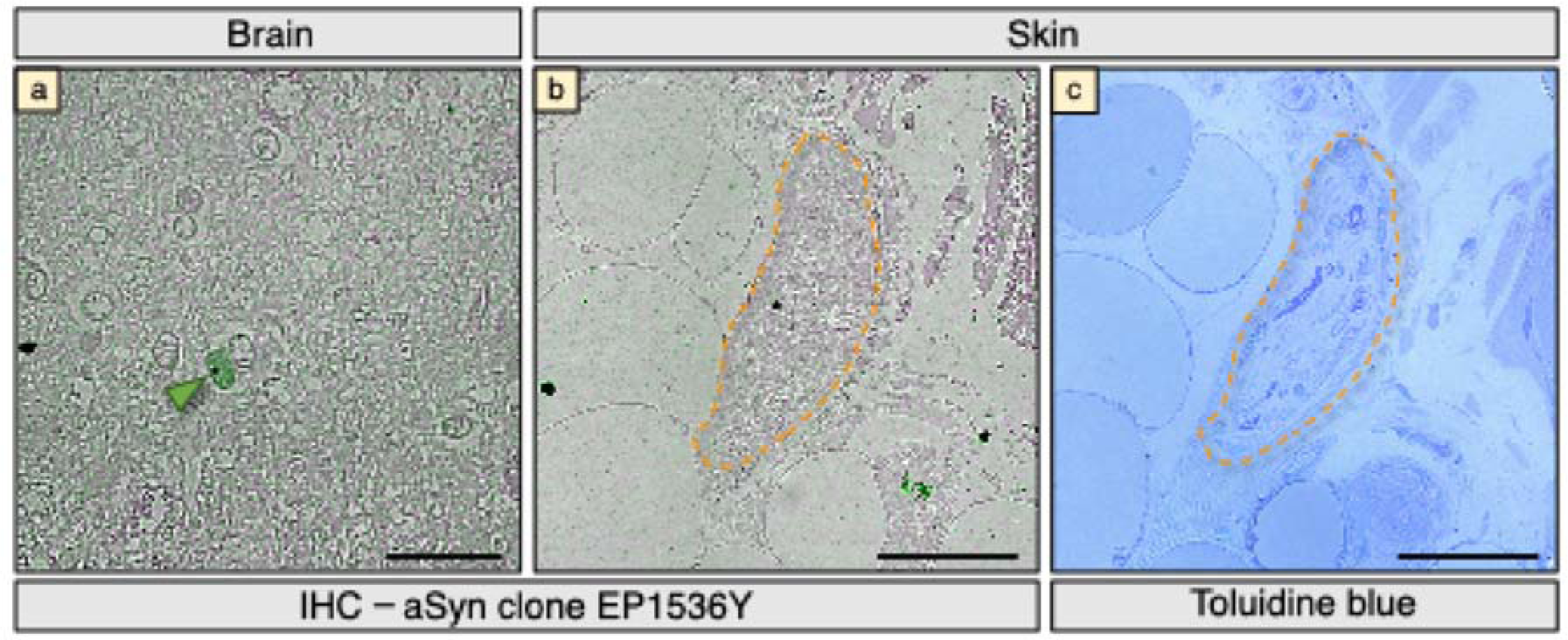
Immunohistochemistry against aSyn in 200 nm brain and skin sections. (a) Example of immunohistochemistry performed against aSyn clone EP1536Y primary antibodies in the brain (*substantia nigra*) of a MSA donor. Arrowhead: glial cytoplasmic inclusions positive for aSyn. (b) Example of immunohistochemistry against aSyn with the same antibody used in the brain. No staining is visible in the skin nerve fiber bundle. Orange dashed lines: myelinated nerve fiber bundle. (c) Consecutive section stained with toluidine blue to highlight skin tissue morphology and locate myelinated nerve fiber bundle shown in (b). Immunohistochemistry experiments performed with the other seven aSyn antibodies listed in Supplementary Table 2 are not shown but produced similar results. Scale bars a, b: 20 μm; c-f: 50 μm.

## Discussion

In this study we used CLEM to characterize the ultrastructure of myelin sheaths at nanometer resolution in human dermal nerve fiber bundles from post-mortem donors diseased with synucleinopathies (PD, DLB, MSA) and non-neurological controls. We combined this technique with ELISA to quantify levels of MPZ, a key protein responsible for myelin sheath compaction, to determine whether myelin damage correlates with changes in myelin protein levels and assess its potential as a disease biomarker.

We observed swellings and separation in myelin sheaths that we classified in four damage categories ranging from absent to high damage. We report a significant difference in the load of myelin damage in our cohort, with cervical skin from PD donors showing significantly higher myelin damage compared to control and MSA groups. We defined a total myelin damage score that can be used as a disease classifier, showing 75% sensitivity, 87% specificity, 81% accuracy, 86% PPV and 78% NPV, suggesting an overall good performance of this classification. Additionally, we report the AUC^45,46^, a key measure of diagnostic accuracy as 0.79 and 0.91 for CTRL-PD and MSA-PD respectively. Since an AUC > 0.8 is generally considered to be clinically useful, our scores indicate that there is a 79% and 91% probability of correctly distinguishing a PD donor from a control or MSA donor.

Notably, we found no significant correlation between myelin damage and age at death, post-mortem delay or fixation time. Additionally, ANCOVA analysis showed no influence of age in myelin damage, thickness and g-ratio. This suggests that, in our cohort, these variables did not significantly impact the differences in myelin damage across donor groups. Taken together, our results support the hypothesis that synucleinopathies may differentially affect peripheral innervation, contributing to the observed variations in myelin damage.

Examining myelin damage in the skin did not allow us to distinguish between the PD and DLB groups. These results support the classification of DLB as “one of the phenotypes in the broader spectrum of Lewy body disease, highlighting its similarities with PD^47–50^. Despite this, we observed pathological differences in the brain tissue, with varying stages of amyloid beta and tau pathology across our cohort (Supplementary Table 1). However, the small sample size of the DLB group and the heterogeneity of myelin damage in this subset limit the ability to draw definitive conclusions.

Conversely, we observed lower levels of myelin damage in the MSA group compared to the PD group. This discrepancy may be partially explained by the donor age, as MSA donors were significantly younger than the other groups due to earlier symptom onset and shorter disease duration. However, despite age differences between the MSA and control groups, the PD group still showed significantly higher myelin damage. This suggests that factors beyond age, such as disease-specific mechanisms, likely contribute to the extent of myelin damage observed in PD.

Previous studies report that up to 55% of PD patients exhibit peripheral neuropathy^33,34^ but it remains unclear whether this is caused by the disease itself or the effects of medications such as levodopa administration. Signs of peripheral neuropathy are present in 9% of the normal population^34^ and levodopa-naïve^31,51^ PD patients. Since MSA patients typically respond poorly to levodopa^4^, differences in levodopa exposure could partly explain the different levels of nerve fiber preservation we observed between Lewy body and MSA donors. However, we did not find a significant correlation between levodopa daily equivalent dose and myelin damage, suggesting that the observed pathology is independent of levodopa treatment.

Another possible explanation for the different levels of myelin damage between PD and MSA donors could be linked to a differential and selective vulnerability of nerve fibers in the two groups^52,53^. Many observations show that cutaneous aSyn depositions in PD and DLB are mainly found in autonomic fibers, while in MSA, they predominantly affect somatosensory fibers^25,26,28,54,55^. In our study, we could not differentiate between autonomic or somatosensory fibers based on morphology alone, and there are currently no antibodies available that we could use for CLEM to distinguish between these fiber types. Further studies are needed to explore this selective vulnerability in greater detail.

The differential biopsy site also seems to affect the presence of aSyn inclusions in the skin: a proximal to distal gradient is reported for PD and DLB^25,26,28^, while a distal to proximal gradient^25,26,28^, or a widespread^54^ distribution is reported for MSA. Since all skin biopsies in this study were collected from the cervical site, our findings may reflect these patterns reported in the literature, where proximal nerve fiber bundles might be impaired in PD and DLB due to the aSyn deposition, but are relatively preserved in MSA. A similar study conducted on distal biopsy sites could provide more precise insights into this hypothesis.

We also stained a subset of myelinated fibers with a set of aSyn antibodies and found that the examined nerve fiber bundles were locally negative for aSyn. This result is consistent with our TEM observations, where no obvious aSyn inclusions were observed in the myelinated nerve fiber bundles, and no pathologies typically found in the brain of individuals affected by synucleinopathies were present. Previous studies have reported the presence of aSyn deposits in the dermal nerve fibers of subjects affected by synucleinopathies^24–31^. Using confocal microscopy on 6 µm thick FFPE sections, we detected pS129 in both myelinated and non-myelinated axons (Supplementary Figure 7). However, in our images as well as in previous reports, aSyn staining was found to be highly localized to specific sites in the nerve fibers, making imaging challenging within the 200 nm thin sections used for CLEM in this study. Consequently, we were not able to definitively determine if the observed damage in the nerve fiber bundles correlates with aSyn accumulation in the same fibers. If myelin damage and aSyn presence were related, it could suggest a Wallerian degeneration-like process^56,57^, where aSyn may trigger neuronal damage or dysfunction in the soma or in the axon, leading to distal degeneration^48^.

We hypothesize that the observed alterations in myelin structure in the skin tissue of PD cases could be due to changes in myelin protein content. To investigate this, we developed and validated an in-house ELISA to measure the amount of MPZ in skin tissue homogenates. However, the ELISA results revealed no differences between PD, DLB, MSA and control cases. This suggests that the structural integrity of myelin proteins may be altered instead, affecting their functionality in compacting the myelin sheaths. Alternatively, changes in the levels of other myelin proteins, such as myelin basic protein, or a biophysical alteration of the lipidic membrane state, could contribute to the altered myelin compaction.

Finally, we showed that myelin damage positively correlates with the presence of hallucinations, as well as LB brain pathology. Further studies will be needed to clarify, if myelin damage in the cervical skin is a specific feature of PD, or if it is also present in other diseases, in absence of aSyn pathology, such as Alzheimer’s disease. Hallucinations are present in many neurological disorders such as PD, DLB and schizophrenia^58^. Notably, previous magnetic resonance imaging (MRI) studies have reported an association between hallucinations and white matter tract abnormalities^59–61^ in the central nervous system in both schizophrenia and PD, supporting a possible link between myelin integrity and hallucination-related circuitry.

This study has two main limitations. First, the relatively small sample size, especially for the rarer synucleinopathies (DLB and MSA) limits the generalization of our findings. Collecting post-mortem samples with such a high level of tissue preservation is extremely challenging, and only few brain banks in the world can collect tissue with the short post-mortem delay required for TEM ultrastructural studies. However, this limitation is mitigated by the high number of myelin sheaths we analyzed with TEM (1123 in total, 104 and 170 from DLB and MSA donors, respectively), which allowed for a robust analysis. Second, as our cohort is post-mortem, we were limited to end-stage diseased subjects and were unable to conduct any follow-up studies. However, this limitation is also one of the main strengths of our study, as it enables precise pathological assessment that is often not possible with living cohorts due to clinical-pathological mismatches, often complicated by concomitant pathology^48^. Therefore, analyzing a post-mortem cohort allowed us to unequivocally assess disease-specific differences.

The abnormalities in the myelin sheaths surrounding nerve fibers that we report in this study can help to discriminate between synucleinopathies and to understand the involvement of peripheral innervation in the diseases. This study should now be extended to living patients to investigate myelin disturbances in healthy controls, different stages of synucleinopathies, and non-synuclein diseases. This would assess whether myelin damage in the peripheral nervous system can be used as a biomarker for disease diagnosis and differentiation, and if so, whether the level of myelin damage correlates with specific disease stages.

In this study, we observed a correlation between myelin damage and age at death for the MSA group. However, in our study the MSA cohort was quite small (N=5), limiting the statistical power of the test. Additionally, the MSA group was younger than control, PD and DLB cohorts (Supplementary Table 1). Therefore, performing a similar study in living patients would benefit from including age-matched controls for the MSA cohort to better account for age-related changes in myelin structure and function.

The identification of a novel biomarker that is effective in diagnosing and differentiating between the synucleinopathies would have significant implications for early detection, disease monitoring, earlier interventions for future treatments, and better outcomes for the patients who suffer from these debilitating diseases.

## Methods

### Human postmortem brain and skin samples

We included a total number of 46 skin donors (23 PD, 7 DLB, 5 MSA and 11 non-neurological controls). 45 of them participated in the brain and skin donation program from the Normal Ageing Brain Collection Amsterdam (NABCA: www.nabca.eu) and Netherlands Brain Bank (www.brainbank.nl). One donor was enrolled by the UW Biorepository and Integrated Neuropathology (BRaIN) Laboratory in the Department of Laboratory Medicine and Pathology of the University of Washington, Seattle (US). Cervical C6-T1 3-mm skin biopsies were collected with rapid autopsy according to a standardized procedure established by NABCA. Tissue blocks were fixed in 4 % paraformaldehyde and 0.1 % glutaraldehyde in 0.1 M cacodylate buffer, supplemented with 1 mM calcium chloride, pH 7.4 for 18-24 hours in most cases (Supplementary Tables 1 and 3 for details) if used for TEM studies (with a post-mortem delay < 10 hrs), frozen, if used for ELISA quantification or formalin fixed and paraffin embedded (FFPE) if used for immunochemistry on paraffin sections. Groups did not differ for fixation time, PMD and age at death, except for the MSA group that was younger than all the other groups (Supplementary Tables 1 and 3).

All donors provided written informed consent for a brain and skin autopsy and the use of the material and clinical information for research purposes. Detailed neuropathological and clinical information was made available, in compliance with local ethical and legal guidelines, and all protocols were approved by Vrije University Medical Center institutional review board. Demographic features and clinical symptoms were abstracted from the clinical files, including sex, age at symptom onset, age at death, disease duration, presence of dementia, core and supportive clinical features for synucleinopathies.

For pathological diagnosis, seven µm-thick FFPE sections were immuno-stained using antibodies against αSyn (clone KM51, 1:500, Monosan Xtra, The Netherlands), amyloid-β (clone 4G8, 1:8000, Biolegend, USA) and phosphorylated tau (p-tau, clone AT8, 1:500, Thermo Fisher Scientific, USA), as previously described 58. Braak and McKeith αSyn stages were determined using the BrainNet Europe (BNE) criteria^62^. Based on Thal amyloid-β phases scored on the medial temporal lobe^63^, Braak neurofibrillary stages^62^ and CERAD neuritic plaque scores^64^ levels of AD pathology were determined according to NIA-AA consensus criteria^65^. Additionally, Thal CAA stages^66^, presence of aging-related tau astrogliopathy (ARTAG)^67^, microvascular lesions and hippocampal sclerosis were assessed.

### Immunohistochemistry on paraffin sections

For detecting phosphorylated Serine 129 (pS129) aSyn in skin biopsies of synucleinopathy cases, we performed immunohistochemistry. We examined whether pS129 aSyn could be observed in myelinated nerve fibers by staining for myelin protein zero (MPZ) and PGP9.5, a pan-neuronal marker. Firstly, FFPE skin biopsies were cut in 6 µm sections and sections from two distinct biopsies were placed on the same glass slide. Sections were deparaffinized by incubation for 3 x 10 min in 100% xylene, followed by incubation in 100% ethanol (2 × 5 min), 96% ethanol (5 min), 80% ethanol (5 min), 70% ethanol (5 min) after which the section were rehydrated in demi water for 5 min. Antigen retrieval was then performed by steam cooking the sections in 10 mM citrate buffer pH 6.0, after which sections were cooled off to room temp before proceeding. Sections were washed for 5 min in TBS, endogenous peroxidase was blocked with a 3% H2O2 solution for 30 min. Following blocking, sections were washed 3 x 5 min in TBS and a non-specific antibody binding blocking step was performed by incubating the sections in a TBS based 5% NGS with 0.3% Triton X-100 solution for 1 hour at room temperature. Subsequently, primary antibody incubation with a mouse anti-pS129 aSyn antibody (825701 clone p-syn/81A, BioLegend, 1:500 working dilution), rabbit anti-PGP9.5 antibody (ab108986 clone EPR4118, Abcam, 1:500 working dilution) and chicken anti-MPZ antibody (LS-C149138 polyclonal, LSBio, 1:100 working dilution) took place overnight at 4°C. The next day, sections were washed 3 x 5 min in TBS, after which secondary antibody incubation took place. Secondary antibody incubation using goat anti-mouse STAR580 (Abberior, cat# ST580-1001, 1:200 working dilution), goat anti-rabbit STAR635P (Abberior, cat#ST635P-1002, 1:200 working dilution) and goat anti-chicken STAR488 (Abberior, cat#ST488-1005, 1:200 working dilution) took place for 2 hours at room temperature. DAPI was used to visualize cell nuclei. After secondary antibody incubation, sections were washed for 3 x 5 min. Hereafter, sections were mounted Sections were then washed for 3 x 5 min and slides were cover slipped with Mowiol + DABCO as mounting medium.

### Frozen skin tissue homogenization

For the biochemical analysis of MPZ protein levels in skin tissue, we homogenized snap frozen skin biopsies. Skin tissue biopsies were thawed on ice and washed 3 times with ice cold PBS pH 7.4, after which skin biopsies were transferred to a cold surface and finely minced with two razor blades. After mincing, the tissue was transferred to a soft tissue CK14 homogenization tube (Bertin Technologies) containing 1.4 mm ceramic beads. For each biopsy, a homogenization buffer containing 150 mM NaCl, 1% Triton X-100m 5 mM EDTA and protease and phosphatase inhibitors in PBS was added in a 1:20 weight to volume ratio to each tube. Mechanical homogenization was performed using the Precellys tissue homogenizer using the Cryolys cooling attachment. Homogenization was performed for 3 cycles of 30 seconds at 3800 RPM, after which samples were rested at 4[for 5 min and homogenization was performed once more. After the second round of homogenization, tissue homogenates were transferred to Eppendorf tubes and spun down at 10.000 RPM for 3 min. The supernatant was collected, aliquoted and stored at -80[until further use.

### MPZ ELISA development and assay validation

For measuring the amount of MPZ in the skin tissue homogenates, we developed an indirect MPZ ELISA for quantification. Recombinant MPZ (ab114281, Abcam) was prepared as a standard in a 3-fold dilution ranging from 30 ng/mL to 41 pg/mL in a 100 mM bicarbonate buffer pH 9.6 and wells were coated with 100 µL per well. Spiking retrieval and linearity of sample dilution were used as validation for the assay. Samples were incubated at 1:400 dilution in bicarbonate buffer pH 9.6 and the plate was incubated overnight at 4°C. After overnight incubation, plates were washed 3 times with TBS-Tween 0.05% (TBS-T). The plate was then blocked with 5% Normal Donkey Serum (NDS) in TBS-T for 1 hour at 37°C. After blocking, the plate was washed like previously described. MPZ antibody (ab183868 clone EPR20383, Abcam) was prepared at a concentration of 0.1 µg/mL in TBS-T supplemented with 1% NDS and the plate was incubated with the primary antibody solution for 1 hour at 37°C. After incubation, the plate was washed as previously described. After washing, the plate was incubated with a biotinylated secondary antibody solution at dilution of 1:20.000 for 1 hour at 37°C. Hereafter, the plate was washed as previously described. The plate was then incubated with streptavidin poly-HRP (M2032, Sanquin) prepared in TBS-T and incubation took place for 1 hour at 37°C. The plate was then again washed as previously described. 1-Step™ Ultra-TMB substrate (34028, ThermoFisher) was added for color development and incubation took place at room temperature until sufficient color had developed. The reaction was then stopped by adding sulfuric acid to each well and absorbance was measured at 450 nm using a plate reader. Dilution linearity of the sample was determined by diluting the sample 1:400, 1:1600 and 1:6400 by determining the retrieval of the analyte in increasing dilutions. 1:400 was used to compare the calculated and measured concentrations of the analyte in the 1:1600 and 1:6400 dilutions. Percentage recovery was determined for a total of 5 cases. Spiking retrieval was assessed by spiking a known concentration of recombinant MPZ into a sample with high endogenous MPZ levels as was determined in the dilution linearity experiment. The sample without spike was compared to samples with a 370 pg/mL (low spike), 1.11 ng/mL (moderate spike) and 3.33 ng/mL (high spike) concentration. Spiking was performed in a 1:400, 1:1600 and 1:6400 dilution. Sigmoidal, 4PL, X is concentration standard curve was fitted to determine protein concentrations and limit of detection (LOD) using Prism Graphpad. Comparisons between percentage recovery for the dilutions and spike recovery was performed by using the one-way ANOVA with Tukey’s multiple comparisons test, with significance considered to be at p < 0.05.

### Resin embedding

3-mm skin biopsies were manually cut into 1-mm blocks containing epidermis and dermis with double-sided razor blades (E72000, Science Services) and prepared for electron microscopy as described previously^68^. Briefly, the biopsies were post-fixed for 1h in 1 % osmium tetroxide reduced with 1.5 % potassium ferrocyanide in 0.3 M cacodylate buffer, supplemented with 3 mM calcium chloride and later for 1h in 1% osmium tetroxide in 0.3 M cacodylate buffer, supplemented with 3 mM calcium chloride. They were washed in water and later stained with 1 % uranyl acetate for 1h, washed in water and dehydrated in a graded acetone series and en-bloc embedded in Epon resin (EMBED 812 embedding kit with BDMA, 14121, EMS). The resin blocks were cured in the oven at 65 °C for 65 hours.

### Serial sectioning

Serial sections of 80 and 200 nm were cut using an ultramicrotome (Leica UC7) and alternatingly collected on TEM grids and glass slides, respectively. Prior to the collection, 200 nm resin sections were stained with toluidine blue (1% with 1% borax in water) for 1-2 minutes at 90 °C to check the presence of myelinated nerve fiber bundles.

### Immunohistochemistry on resin sections

Some sections on the glass slides were also immunolabeled against MPZ to validate the toluidine blue staining and against aSyn to check the local presence of aSyn inclusions. The sections were etched in a saturated potassium ethoxide solution for 3 mins followed by washing in ethanol and water. Depending on the primary antibody used, antigen retrieval was carried out if necessary, with 100 % formic acid for 10 minutes followed by steaming in Tris-EDTA, pH 9 or sodium citrate, pH 6, for 30 mins at 100 °C (Supplementary Table 2). Endogenous peroxidases were quenched with 1 % hydrogen peroxide in 10 % methanol, before blocking in antibody diluent (Dako REAL, Agilent). The sections were incubated with primary antibodies overnight at 4°C, at room temperature for 4 hours or at 37 °C for 1 hour, depending on the antibody (Supplementary Table 2). The sections were then washed in PBS supplemented with 0.25 % Triton X-100 and incubated in the secondary antibody (ImmPRESS Reagent Anti-Rabbit Ig or anti-mouse, Vector Laboratories) for 30 mins at room temperature. Bound antibody complexes were detected using the permanent HRP Green Kit (Zytomed Systems) with incubation for 3 mins at room temperature. Sections were counterstained with hematoxylin, dehydrated and mounted with glass coverslips for imaging.

### Light microscopy - resin sections

Light microscopy images of selected glass slides containing myelinated nerve fiber bundles (detected either with toluidine blue staining or immunohistochemistry) were collected using a Leica Thunder microscope equipped with a Flexacam color camera. The regions of interest containing myelinated nerve fiber bundles were imaged in overlapping tiles at 630 X (oil immersion) magnification and merged into a single image using the LAS X software (Leica Microsystems). These images were used to find the exact location of the corresponding region of interest on the adjacent TEM grid.

### Electron microscopy

TEM images of electron microscopy grids consecutive to those imaged by light microscopy were collected at room temperature on a 120 kV Tecnai G2 Spirit TEM microscope operated at 80 kV with a LaB6 filament and a bottom mount TVIPS F416 camera, or a CM100 Biotwin (Philips) operated at 80 kV with a Lab6 filament and bottom mount TVIPS F416 camera. Tiled images at 6500 X magnification and pixel size of 3.16 nm were acquired.

### Myelin sheath damage analysis

We analyzed 1123 myelin sheaths with TEM and defined 4 categories of myelin damage (absent N-, mild N+, moderate N++ and high N+++ damage) based on the size and density of myelin sheath swellings visible with TEM. N-includes all the myelinated fibers that do not show any swelling, hole or separation, and show no change in the periodicity of the intraperiod lines (Figure 2a, e and Supplementary Figure 1a-e). N+ is defined by sparse swellings, while the periodicity of the myelin sheaths is still clearly visible in most of the myelinated fiber (Figure 2b, f and Supplementary Figure 1f-j). Myelinated fibers classified as N++ show moderately frequent swellings and the periodicity of the myelin sheaths is still visible in some segments of the myelinated fiber but is heavily disrupted in many others (Figure 2c, g and Supplementary Figure 2a-e). Finally, N+++ is defined by highly frequent swellings and the periodicity of the myelin sheaths is poorly visible and heavily disrupted in most of the myelinated fiber (Figure 2d, h and Supplementary Figure 2f-j). We defined a total score for each donor as a weighted mean of the 4 categories of myelin damage (*Score_tot_* 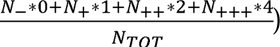 giving different weights (0, 1, 2, 4) to increasing levels of damage. *N*_-_, *N*_+_, *N*_++_, *N*_+++_ are the number of myelin sheaths with no, mild, moderate, or high damage, respectively, and *N_TOT_* is the sum of all of them. The total score was used to show differences among disease groups and as a disease classifier. We did not include Schmidt-Lanterman incisures and nodes of Ranvier in the count of myelin damage.

### Myelin sheath thickness and g-ratio analysis

Myelin sheaths and axons were segmented in Fiji^69^ using the semi-automated SAMJ-IJ plugin (https://github.com/segment-anything-models-java/SAMJ-IJ) with manual prompts and manually refined where needed. Custom Fiji macros were used to automate myelin label saving and myelin thickness calculation using the Fiji built-in function Local Thickness (complete process). Average thickness values were automatically extracted from myelin sheaths with custom routines in Matlab R2023a. The g-ratio was calculated in Matlab R2023a from myelin sheaths from each donor processed for EM using the formula^70^: 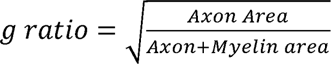 as the axon shape was often irregular. EM myelin sheath images with tissue folds were excluded for the calculation of the g-ratio. Additionally, images > 10MB were above the size limit for the Local Thickness function in Fiji and so were also excluded.

### Electron tomography

Tilt-series were collected on a 120 kV Tecnai G2 Spirit TEM microscope operated at 120 kV equipped with a LaB6 filament and a bottom mount TVIPS F416. Images were collected in 1° steps from -60° to +60° sample tilt using the SerialEM software^71^ and with a pixel size of 0.43 nm. Tomograms were binned by a factor of 2 and filtered using a non-local means filter in Amira version 2021.2 (ThermoFisher Scientific). Segmentation of the myelin sheaths was carried out in Fiji and Amira.

### Statistical analysis

Raw data were processed in Microsoft Excel and statistics were performed using Prism Graphpad (version 9.3.1) www.graphpad.com, MATLAB R2023a^72^ and R^73^. Sensitivity, specificity, positive predictive value, negative predictive value, and accuracy were calculated using the pathological diagnosis as the gold standard. Shapiro-Wilk test was used to verify the normal distribution of continuous variables. ANCOVA-analysis was performed to evaluate the effect of categorical variables (here, disease group) on a continuous dependent variable (total myelin damage score), while adjusting for potential confounders (age). Two sample t-test was used to compare two groups with normal distribution. A Chi-squared test was used to statistically assess the distribution of myelin damage across the disease groups. Kruskal-Wallis test followed by Dunn’s test were utilized to compare variables without a normal distribution. Resulting p-values were corrected for multiple comparisons according to the Bonferroni method. Pearson or Spearman correlation coefficients were calculated according to the variable type and distribution. The p-values in the correlation matrix were adjusted with the Benjamini-Hochberg procedure. For all analyses, significance was assumed for corrected p-values < 0.05. Empty circles in box plots are outliers calculated with the standard interquartile range method.

## Abbreviations

aSyn: alpha synuclein
CLEM: correlative light and electron microscopy
DLB: Dementia with Lewy bodies
ELISA: enzyme linked immunosorbent assay
IHC: immunohistochemistry
LB: Lewy body
MSA: multiple system atrophy
MPZ: myelin protein zero
PD: Parkinson’s disease
TEM: transmission electron microscopy

## Data availability

All data will be made available on request.

## Code availability

Custom codes used in this study are available at https://github.com/LBEM-CH/myelino

## Acknowledgements

We would like to thank the donors and their families who participated in the brain donation program to make this study possible. We would like to thank all members of the Netherlands Brain bank and the BRaIN lab autopsy teams for facilitating the collection of high-quality postmortem brain tissue for electron microscopy. We thank Roche for providing some of the aSyn antibodies used in this study. We thank C. Genoud, J. Daraspe, D. De Bellis and A. Mucciolo from the Electron Microscopy Facility, University of Lausanne, for training and assistance with tissue processing and electron microscopy. We thank D. Sage (Biomedical Imaging Group, EPFL) and S. Kasas (Laboratory of Biological Electron microscopy, EPFL), for providing knowledge and help with myelin segmentation and SAMJ-IJ. We thank D. Stähli, N. Shafiei, L. van den Heuvel and K. Ekundayo for fruitful discussions.

## Author contributions

MDF, AJL, WvdB and HS designed the study. WvdB collected, dissected and processed the brain samples for pathological diagnosis confirmation. WvdB and BvdG collected the skin biopsies at autopsy. MDF, MT, EA and NbcS performed the CLEM experiments and analysis. MDF and MT performed immunohistochemistry on resin sections. MDF performed the myelin thickness and g-ratio analysis. MDF and NbcS performed EM tomography. MDF performed segmentation of tomograms. BvdG performed the ELISA on frozen biopsies and immunohistochemistry on paraffin sections. MDF and AJL wrote the manuscript with contributions from all authors. All authors contributed to the analysis and interpretation of the data and approved the final version of the manuscript.

