## Supplementary information for "Multiscale analysis of myelin alterations in skin biopsies from synucleinopathies"

**Supplementary Material**

|  | **CTRL** | **PD** | **DLB** | **MSA** |
| --- | --- | --- | --- | --- |
| **Number** | 11 | 24 | 6 | 5 |
| **Sex (F/M)** | 7/4 | 6/18 | 1/5 | 3/2 |
| **Age at death**^a^ **(years ± SD)** | 79.9 ± 9.2 | 76.9 ± 6.1 | 77.0 ± 3.6 | 60.0 ± 7.3 |
| **Age at death**^b^ **(years ± SD)** | 79.9 ± 9.2 | 76.9 ± 6.0 | 77.3 ± 2.9 | 60.0 ± 7.3 |
| **Age at onset (years ± SD)** | na | 60.4 ± 10.2 | 70.2 ± 5.3 | 54.6 ± 9.6 |
| **Disease duration (years ± SD)** | na | 16.4 ± 7.6 | 7.2 ± 3.8 | 5.2 ± 2.3 |
| **PMD**^a^ **(h)** | 4.7 ± 1.5 | 4.2 ± 1.2 | 5.0 ± 2.5 | 5.5 ± 2.4 |
| **PMD**^b^ **(h)** | 4.7 ± 1.5 | 4.9 ± 2.0 | 5.9 ± 2.4 | 5.5 ± 2.4 |
| **Fixation time**^a^ **(h)** | 33 ± 23 | 29 ± 10 | 24 ± 1 | 44 ± 22 |
| **Thal amyloid phase** | 0-4 (4/4/0/2/1*) | 0-4 (5/8/6/3/2) | 0-5 (1/0/0/3/1/1) | 0-4 (3/1/0/0/1) |
| **Braak NFT stage** | 0-3 (1/3/1/6) | 0-3 (5/5/9/5) | 2-4 (1/3/2) | 0-3 (3/1/0/1) |
| **Braak aSyn stage** | 0-6 (9/1/0/0/0/0/1**) | 5-6 (1/23) | 6 (6) | 0-5 (1/0/1/0/0/3) |

**Supplementary Table 1:** Clinicopathological characteristics of donors included in this study. PMD: post-mortem delay, NFT: neuro-fibrillary tangle, LB: Lewy body, ^a^: donors for EM studies, ^b^: full cohort. Brackets indicate the number of donors within each stage. *Non-neurological control with age-associated pathology (#Case 3). **Non-neurological control with age-associated pathology (incidental LB) (#Case 2).

| **Name** | **ID #** | **Company** | **Target** | **Tissue - Brain** | | **Tissue - Skin** | |
| --- | --- | --- | --- | --- | --- | --- | --- |
|  |  |  |  | **AR** | **Condition** | **AR** | **Condition** |
| MPZ – P0 | ab183868 | Abcam | MPZ – aa 1-258 | - | - | yes^a^ | o/n, 4°C, 1:100 |
| LB509 | [180215](https://www.thermofisher.com/order/catalog/product/180215) | Thermofisher | aSyn - aa 115-122 | no | 1h, 37°C, 1:500 | yes^b^ | o/n, 4°C, 1:100 |
| 11A5 | - | Prothena | aSyn - pS129 | no | 1h, 37°C, 1:100000 | yes^b^ | o/n, 4°C, 1:100 |
| aSyn clone EP1536Y | [ab51253](https://www.abcam.com/alpha-synuclein-phospho-s129-antibody-ep1536y-ab51253.html) | Abcam | aSyn - pS129 | no | 1h, 37°C, 1:5000 | yes^b^ | o/n, 4°C, 1:100 |
| aSyn C terminal | [A15127A](https://www.biolegend.com/en-gb/products/purified-anti-alpha-synuclein-c-terminal-truncated-x-122-13986) | Biolegend | aSyn - CTT 122 | yes^b^ | o/n, 4°C, 1:500 | yes^b^ | o/n, 4°C, 1:100 |
| aSyn N terminal | [A15110D](https://www.biolegend.com/en-us/products/purified-anti-alpha-synuclein-34-45-antibody-14067?Clone=A15110D) | Biolegend | aSyn - aa 34-45 | yes^b^ | o/n, 4°C, 1:500 | yes^b^ | o/n, 4°C, 1:100 |
| 131 Roche | - | Roche | aSyn - CTT 119 | no | 1h, 37°C, 1:500 | yes^b^ | o/n, 4°C, 1:100 |
| 134 Roche | - | Roche | aSyn - CTT 122 | no | 1h, 37°C, 1:500 | yes^b^ | o/n, 4°C, 1:100 |
| 142 Roche | - | Roche | aSyn - pS129 | no | 1h, 37°C, 1:500 | yes^b^ | o/n, 4°C, 1:100 |

**Supplementary Table 2: Primary antibodies used in this study for immunohistochemistry on 200 nm resin sections.** ^a^ :10’ in formic acid at RT + 30’ at 95 °C in Tris-EDTA pH9. ^b^ :10’ in formic acid at RT + 30’ at 95 °C in sodium citrate pH6 or 10’ in formic acid at RT + 30’ at 95 °C in Tris-EDTA pH9. AR: antigen retrieval, MPZ: myelin protein zero, aa: amino acids, CTT: C-terminal truncated, o/n: overnight, RT: room temperature.

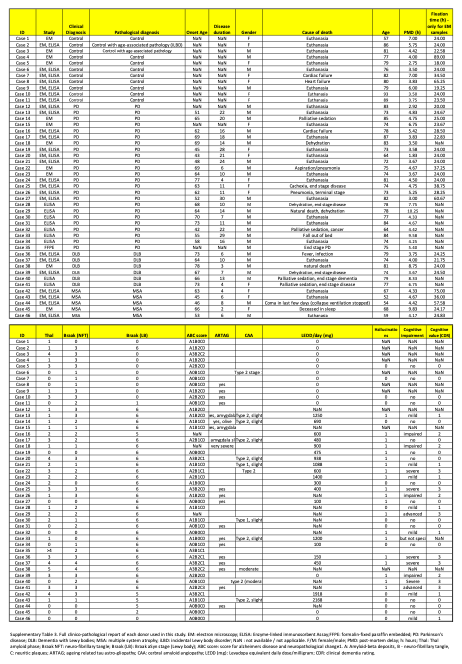

**Supplementary Table 3. Full clinico-pathological report of each donor used in this study.** EM: electron microscopy; ELISA: Enzyme-linked immunosorbent Assay;FFPE: formalin-fixed paraffin embedded; PD: Parkinson's disease; DLB: Dementia with Lewy bodies; MSA: multiple system atrophy; iLBD: incidental Lewy body disorder; NaN: not available / not applicable. F/M: female/male; PMD: post-mortem delay; h: hours; Thal: Thal amyloid phase; Braak NFT: neuro-fibrillary tangle; Braak (LB): Braak aSyn stage (Lewy body); ABC score: score for Alzheimer’s disease and neuropathological change^1,2^. A: Amyloid-beta deposits, B - neuro-fibrillary tangle, C: neuritic plaques; ARTAG; ageing related tau astro-gliopathy; CAA: cerbral amyloid angiopathy; LEDD (mg): Levadopa equivalent daily dose/milligram; CDR: clinical dementia rating.

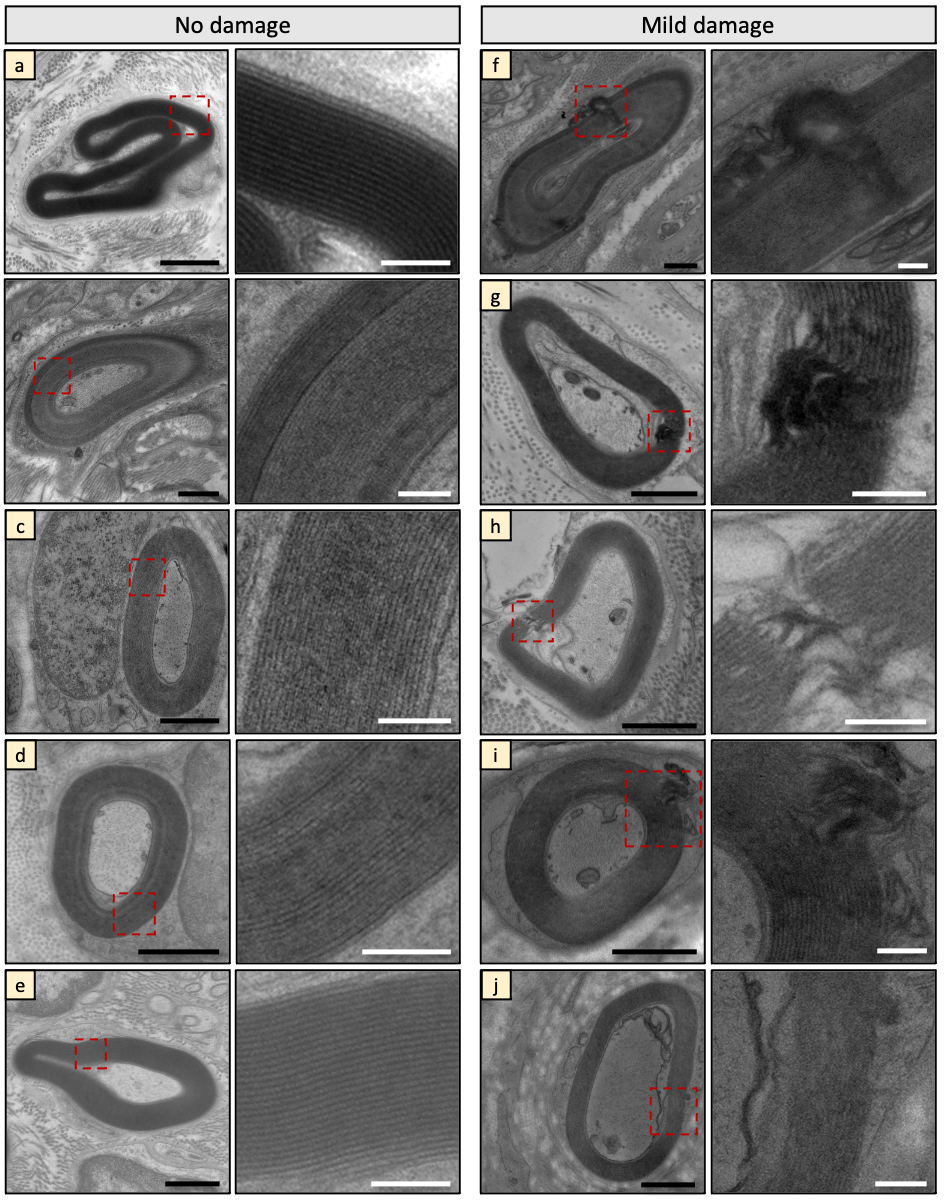

**Supplementary Figure 1: EM images of dermal myelin sheaths (no and mild damage categories).** Examples of non-damaged (a-e) and mildly damaged (f-j) myelinated axons with their zooms (dashed red square). Scale bars: low magnification 1 μm, high magnification 200 nm.

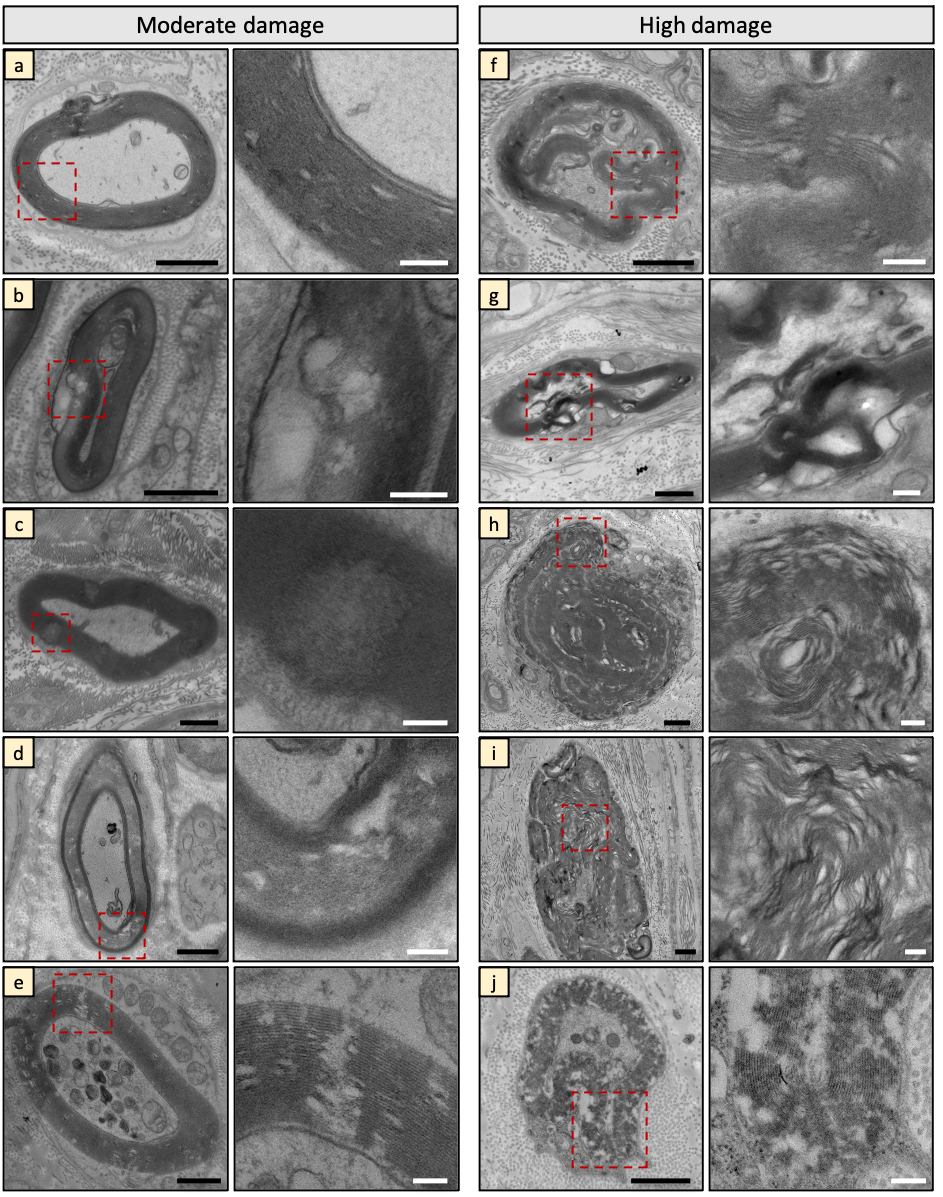

**Supplementary Figure 2: EM images of dermal myelin sheaths (moderate and high damage categories).** Examples of moderately (a-e) and highly damaged (f-j) myelinated axons with their zooms (dashed red square). Scale bars: low magnification 1 μm, high magnification 200 nm.

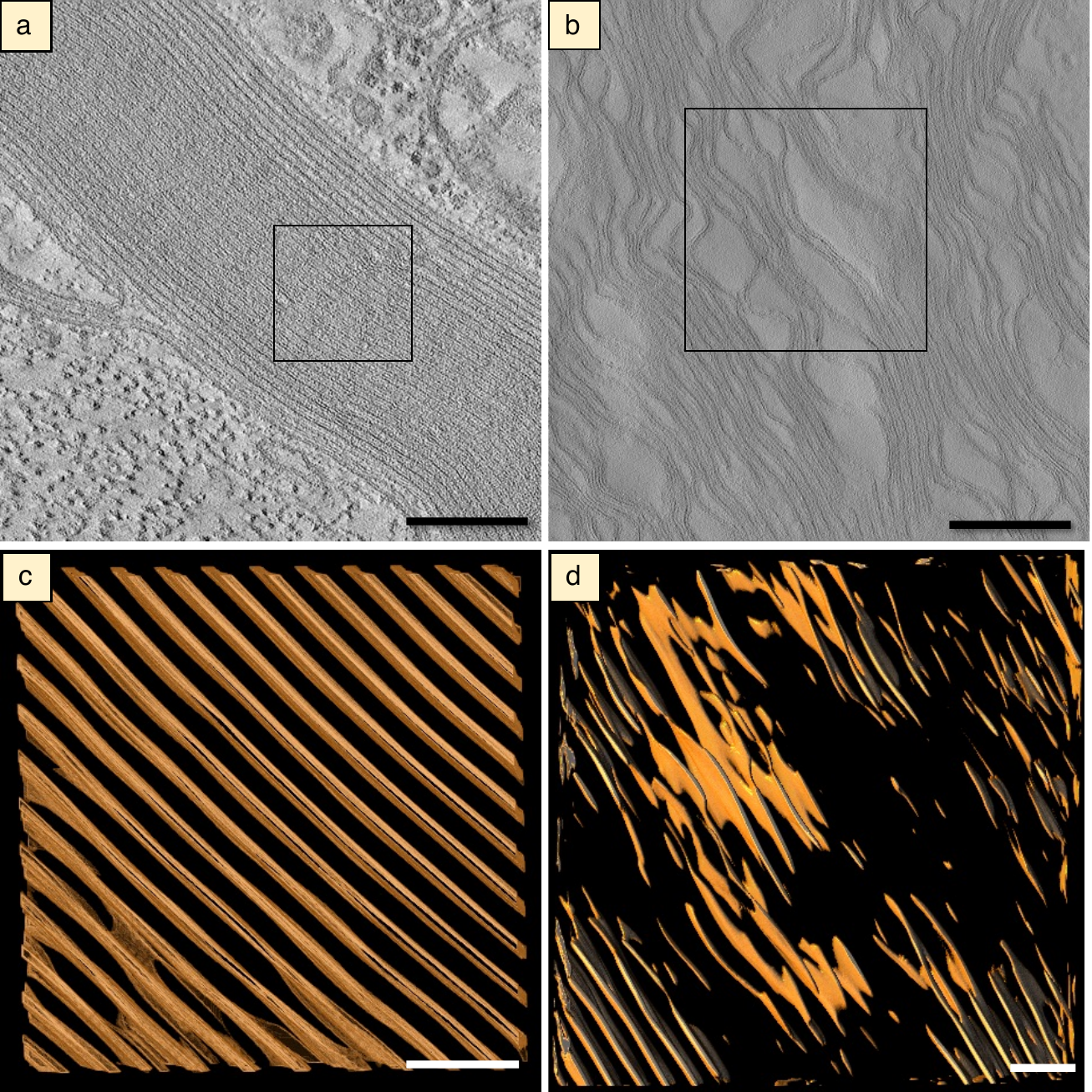

**Supplementary Figure 3: Examples of electron tomography and segmentation of no and high myelin damage categories.** EM tomography from the preserved and highly damaged categories of myelin damage. (a) Tomogram and (b) segmentation of a myelinated axon that did not show any damage. (c) Tomogram and (d) segmentation of a myelinated axon that showed extensive damage. The black boxed in (a) and (b) indicate the segmented area shown in (c) and (d). Swellings within the myelin sheaths are clearly visible in the highly damaged example. The preserved myelin sheath shows repeated and conserved periodicity of the myelin sheaths. Scale bars a,b: 200 nm, c,d: 50 nm.

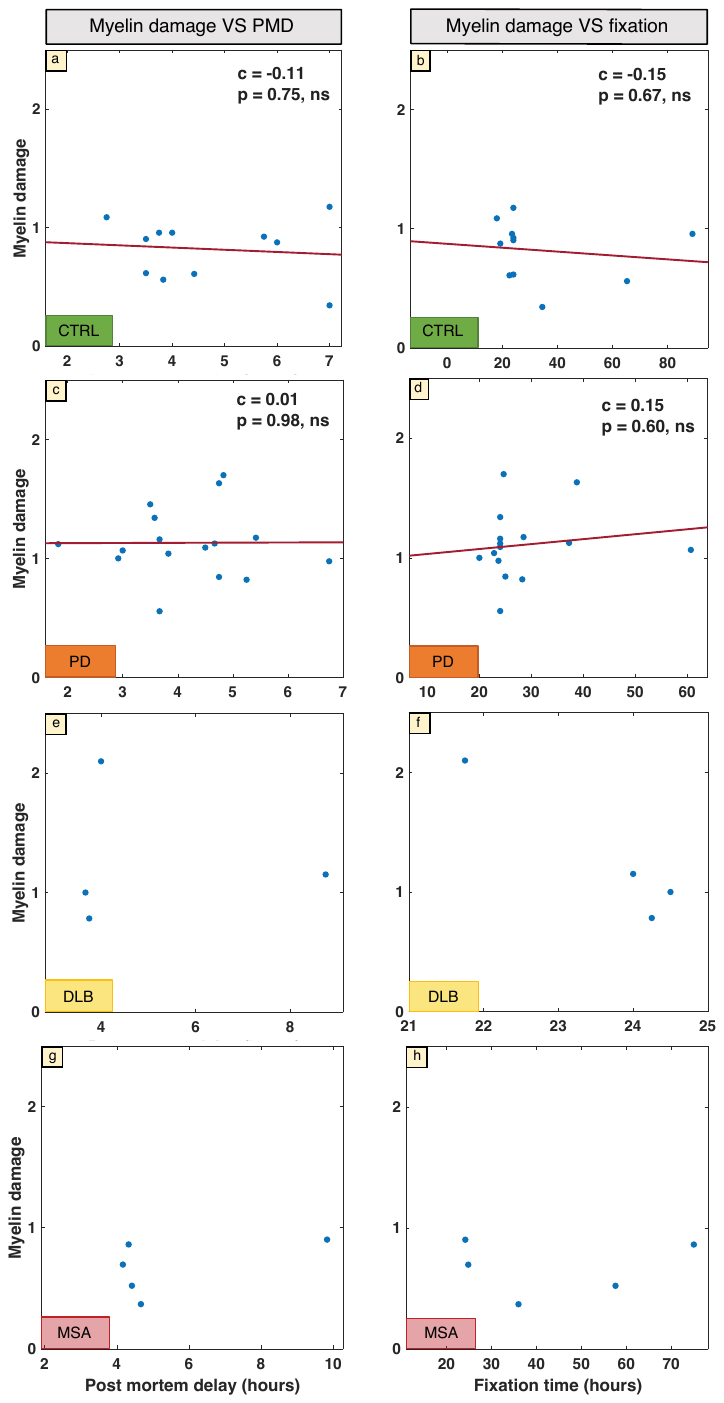

**Supplementary Figure 4: The effect of post-mortem delay (PMD) and fixation time on the presence of myelin damage within the donor groups.** Correlation plots of total myelin damage score as a function of PMD (a-c) and fixation time (b-d) for controls and PD. We did not find any significant correlation between myelin damage and PMD or fixation in our study in the single groups, suggesting they may not be the leading cause of the observed differences in myelin damage. The DLB and MSA groups (e-h) were excluded from the statistical analysis due to the small sample size (N=4 and N=5, respectively). c = Spearman correlation coefficient, ns = non-significant.

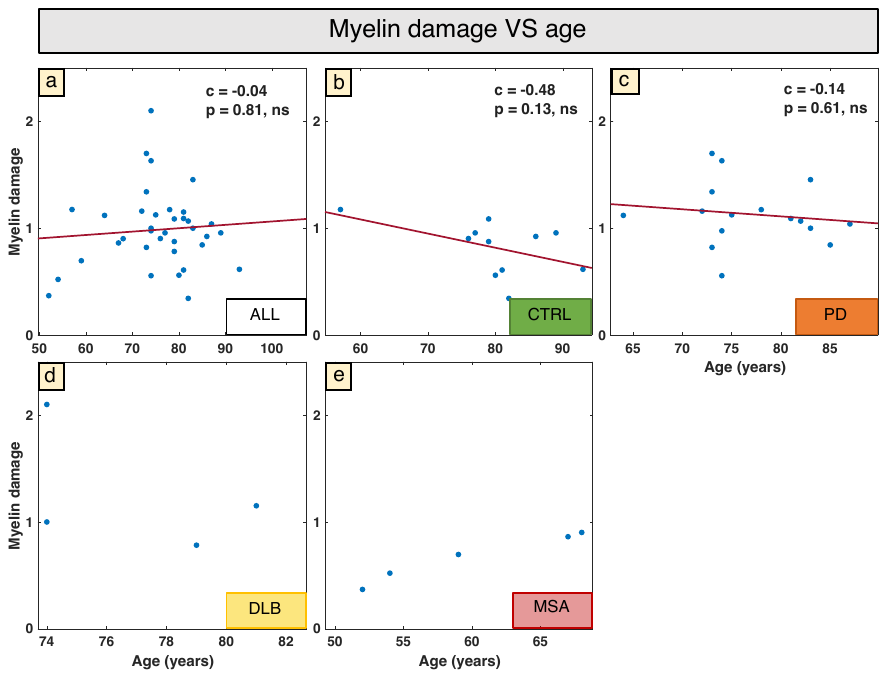

**Supplementary Figure 5: The effect of age at death on the presence of myelin damage in the whole cohort and within the donor groups.** Correlation plots of total myelin damage score as a function of age in the full cohort (a) and in the groups: CTRL (b), PD (c), DLB (d) and MSA (e). No significant correlation between myelin damage and age at death was observed for any group except MSA (c=0.98, p=0.004). However, due to the small sample size in the MSA group (N=5), this correlation should be interpreted with caution, as small samples can produce inflated correlation coefficients. Therefore, both the DLB (N=4) and MSA groups were excluded from analysis. Overall, the level of damage in the MSA and control groups is comparable, despite age differences, while PD and control groups showed differing levels of damage despite comparable age. These factors suggest that age might not be the leading cause of the observed differences in myelin damage. c = Spearman correlation coefficient, ns = non-significant.

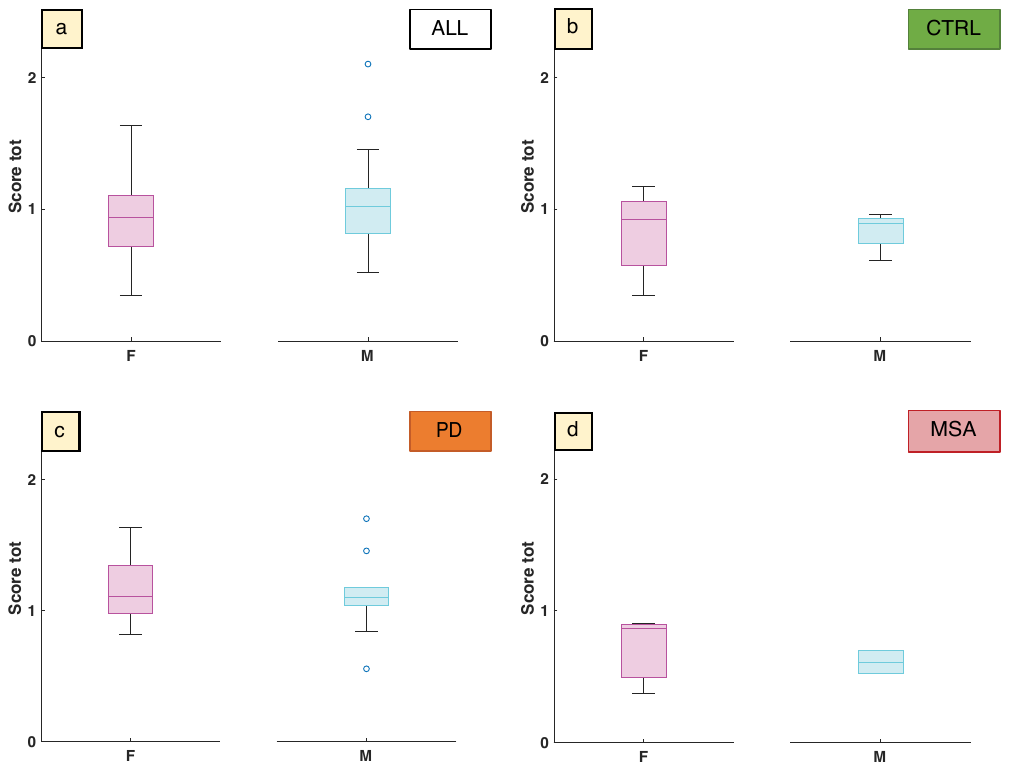

**Supplementary Figure 6: Myelin damage does not show sex differences**. None of the groups show sex differences in terms of myelin damage as all the p-values are non-significant. Total myelin damage score (Score tot) shown for female (pink) and male (blue) groups. (a) full cohort, (b) control, (c) PD and (d) MSA groups. DLB group not shown as no females were included in the EM study.

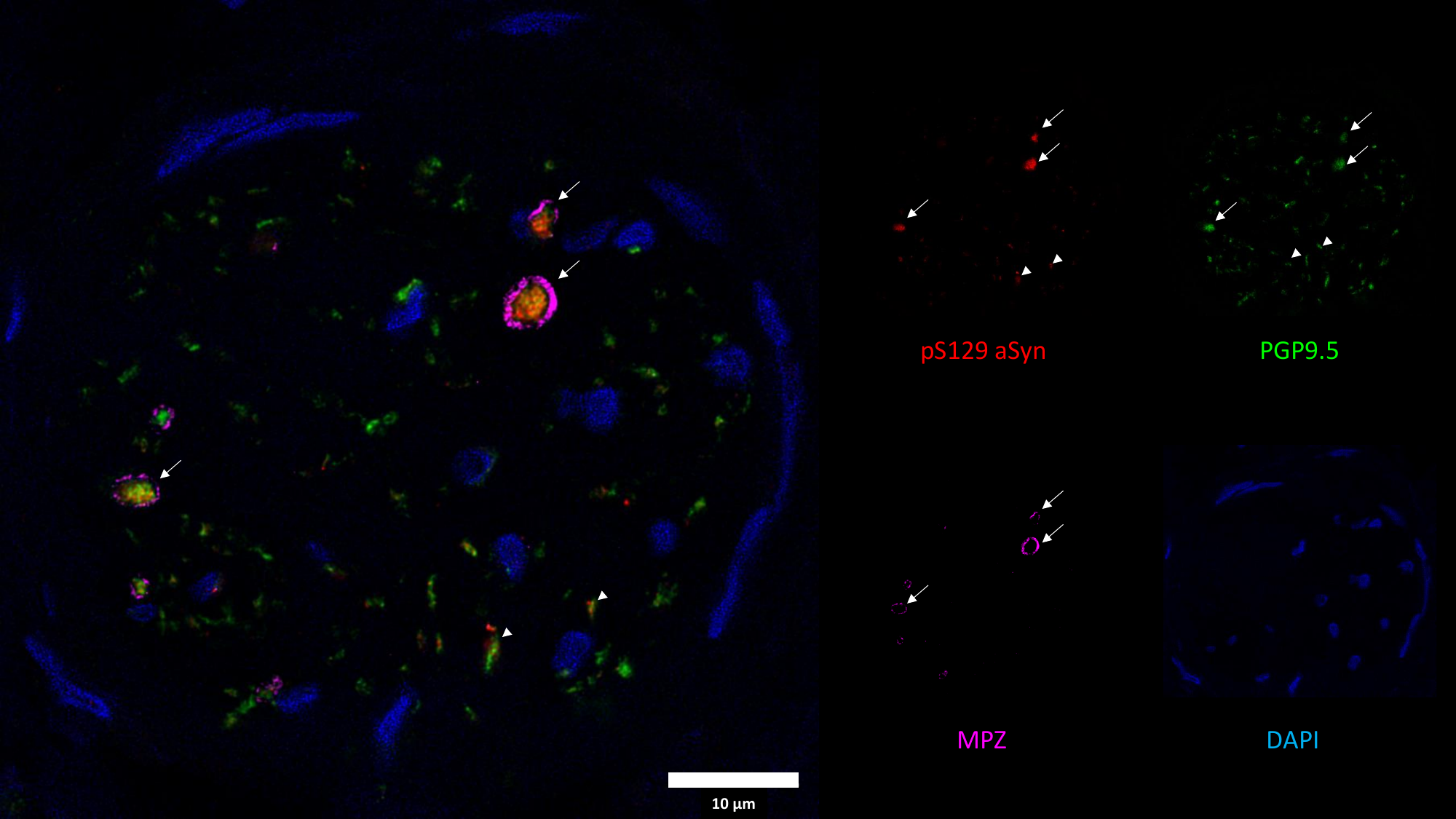

**Supplementary Figure 7: Example of pS129 aSyn deposition observed in myelinated and unmyelinated nerve fibers in a PD patient**. Staining highlighting presence of pS129 aSyn (red) in nerve fibers (green) with (arrow) and without (arrowhead) myelin sheaths (magenta). DAPI (blue) was used to visualize cell nuclei. Scale bars: 10 µm.

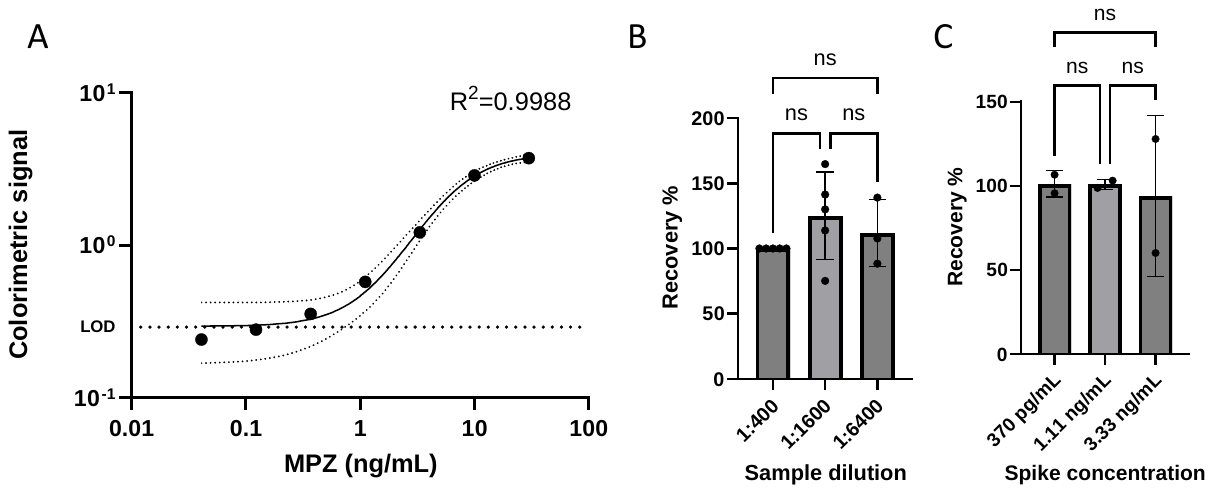

**Supplementary Figure 8. MPZ ELISA validation.** In-house produced indirect MPZ ELISA had a limit of detection of 184 pg/mL, determined by back calculation of the blank plus three times the standard deviation of the blank based on the standard curve (A). Dilution linearity was confirmed by going from 1:400 (set at 100% recovery) to 1:1600 (112% recovery) and 1:6400 (125% recovery) dilutions with no significant differences between any of the dilutions. For the 1:6400 dilution, 2 out of 5 cases fell below the LOD (B). The mean percentage recovery for the spiking was 101% for the low (370 pg/mL) and moderate (1.11 ng/mL) spiking conditions, and 94% recovery for the high (3.33 ng/mL) spiking condition (C) There were no significant differences between any of the conditions, confirming the validity of the assay.
